# A Versatile and Open Source One- and Two-Photon Light-Sheet Microscope Design

**DOI:** 10.1101/2023.07.10.548107

**Authors:** Antoine Hubert, Thomas Panier, Geoffrey Migault, Hugo Trentesaux, Benoît Beaudou, Georges Debrégeas, Volker Bormuth

## Abstract

Two-photon light sheet microscopy offers great potential for a range of biological applications. Still, its practical implementation is impeded by the high cost of laser sources, the complexity of construction, and the challenges associated with adapting to existing microscope setups. Here, we release an open-source design that addresses these limitations by providing detailed building instructions for transforming a brightfield microscope into a versatile one- and two-photon light sheet system. Our design incorporates a specially designed broadband hollow core fiber, enabling the simultaneous utilization of visible laser alongside an expansive pulsed laser source from another setup. This integration allows for uncompromised image resolution and speed. Furthermore, the design reduces the complexity of construction, alignment, and overall cost, thereby significantly enhancing the accessibility of this technology (https://github.com/LJPZebra/OLU).

## Introduction

Light-sheet fluorescence microscopy (LSFM) has emerged as a cornerstone technique in fields ranging from developmental biology (1, 2) to cell biology (3), organoid research (4, 5) and neuroscience (6–9).

In LSFM, the sample is illuminated by a thin light sheet orthogonal to the fluorescence detection axis, thus co-aligning fluorescence stimulation and detection plane. This minimizes photodamage and bleaching relative to epifluorescence or confocal modalities. By combining optical sectioning, rapid volumetric imaging, and low photodamage, LSFM has enabled detailed studies of embryogenesis (such as in Drosophila or Danio rerio) and real-time monitoring of neural dynamics in entire brains of small organisms (10).

In single-photon LSFM, a continuous-wave (CW) laser in the visible range is conventionally employed. Although such illumination is relatively straightforward, it suffers from scattering and absorption in thicker or more opaque specimens. Compared to single-photon systems, two-photon light sheet microscopes offer an increase in penetration depth and a reduction in the background signal by minimizing photon scattering and absorption in tissues (11, 12). Furthermore, the infrared source used in two-photon imaging lies outside the visible spectrum of most animals, thus preventing interference with the visual system (6, 7) during functional imaging.

However, the dissemination of two-photon light-sheet microscopes remains limited due to several factors: (i) they require a dedicated femtosecond laser source, which is costly; (ii) they are difficult to adapt to existing microscopes and thus require a complete custom-made system; and (iii) they are relatively complex to build and challenging to align.

Recent advances in fiber optics are beginning to address these challenges by enabling flexible broadband delivery of laser sources. They allow both visible and pulsed infrared light for one- and two-photon microscopy to be transmitted through the same fiber, with high transmission efficiency and very low pulse dispersion.

Efficient, dispersion-free, and broadband fiber coupling of a two-photon laser source is a long-standing challenge in the field of two-photon microscopy (13) and of fiber optics design (14). The difficulty arises because standard single-mode optical glass fibers are associated with strong linear and nonlinear pulse dispersion, which reduces the two-photon efficiency. Pre-compensation methods are only efficient at low laser powers and are therefore impractical in the context of fast volumetric two-photon imaging, which requires high photonic fluxes.

The development of single-mode photonic bandgap fibers, in which the light travels through an air-filled hollow core, dramatically reduces dispersion and nonlinear effects and achieves transmission ratios greater than 50% (14, 15). These fibers have been successfully used for pulsed laser delivery in the context of multiphoton imaging (16–20). However, as the light-guiding mechanism is based on creating an optical bandgap, these fibers only allow single-wavelength transmission. They are currently only commercially available for laser wavelengths of 800 nm or 1064 nm (NTK photonics). However, with the rapidly growing collections of genetically encoded actuators and sensors, there is a growing demand for broadband fiber delivery that covers the near-infrared and extends into the visible spectrum. Such capability is essential to fully exploit 2P-LSFM across diverse applications and to facilitate laser source sharing between different setups.

Broadband fiber delivery spanning the visible and the near-infrared spectrum is possible with negative curvature hollow-core photonic crystal fibers (NCF), which do not rely on an optical bandgap to confine the laser light to the fiber’s core (21– 23). The simplest cross-sectional geometry of an NCF is based on a ring of touching or non-touching tubes surrounding the core (22). Core diameter, tube diameter, inter-tube distance, and tube wall thickness control the spectral transmission bands, the attenuation level, the quality of higher-order mode suppression, and the sensitivity of the optical properties to bending. Attenuation levels < 0.07 dB/km and bending loss < 0.03 dB/m are reported (24). Due to the minimal lattice structure and the reduced light interaction with the cladding structure, these fibers have very high damage thresholds and can even be used for very high laser energy delivery of up to 100 *µ*J peak power when the core is vacuum pumped to reduce nonlinear effects at these high powers (25). Negative curvature fibers are on the verge of being used in several applications such as laser micromachining and laser surgery. A recent study demonstrated the successful use of a custom HC-NCF with a transmission band of 600 – 830 nm at < 0.3 dB/m attenuation in the design of a handheld two-photon microscopy scanner for human skin autofluorescence (26). This broadband capability opens the door for experimental setups that require single-photon and two-photon illumination paths without separate hardware.

Here, we describe an open-source one- and two-photon light-sheet microscope that capitalizes on the broadband transmission properties of a custom negative curvature hollow-core fiber. We transform a commercial upright electrophysiology microscope into a multiphoton-capable LSFM system by attaching a compact light-sheet forming unit measuring only a few centimeters. The module houses the essential optics for digitally scanned light-sheet generation, enabling the user to align the microscope with minimal manual adjustments. Once assembled, researchers may connect a CW visible laser and an ultrafast near-infrared laser to the fiber input for interchangeable operation in single-photon and two-photon modes. Step-by-step detailed building instructions and blueprints are provided in the open-source online material (https://github.com/LJPZebra/OLU).

We demonstrate the performance of the system by imaging the brain of larval zebrafish expressing the pan-neuronal calcium indicator GCaMP6f. We perform both static anatomical scans and high-speed volumetric recordings of spontaneous neural activity, illustrating the microscope’s capacity for low-phototoxic, high-contrast functional imaging. Overall, this open-source platform has the potential to democratize 2P-LSFM, making advanced imaging techniques more accessible for a broad range of applications in modern biology.

## Results

### A. Transformation of a Commercial Upright Microscope into a One- and Two-Photon Light-Sheet System

We present a versatile open-source one- and two-photon light-sheet microscope. The design features a compact module for light sheet formation (Figure 1) with a small footprint, measuring 6 × 9 × 10 cm^3^ (27) that can be mounted on standard commercial microscopes. We have successfully integrated this light-sheet unit with an electrophysiology microscope from Scientifica (UK), converting it into a dual-mode one- and two-photon light-sheet system (Figure 1a, Video S1). The 3D model can be explored interactively at https://a360.co/3C0TOAm and https://a360.co/434IdMt. Complete building instructions and blueprints are available online (https://github.com/LJPZebra/OLU) and in print format (see Supplement).

**Fig. 1.**
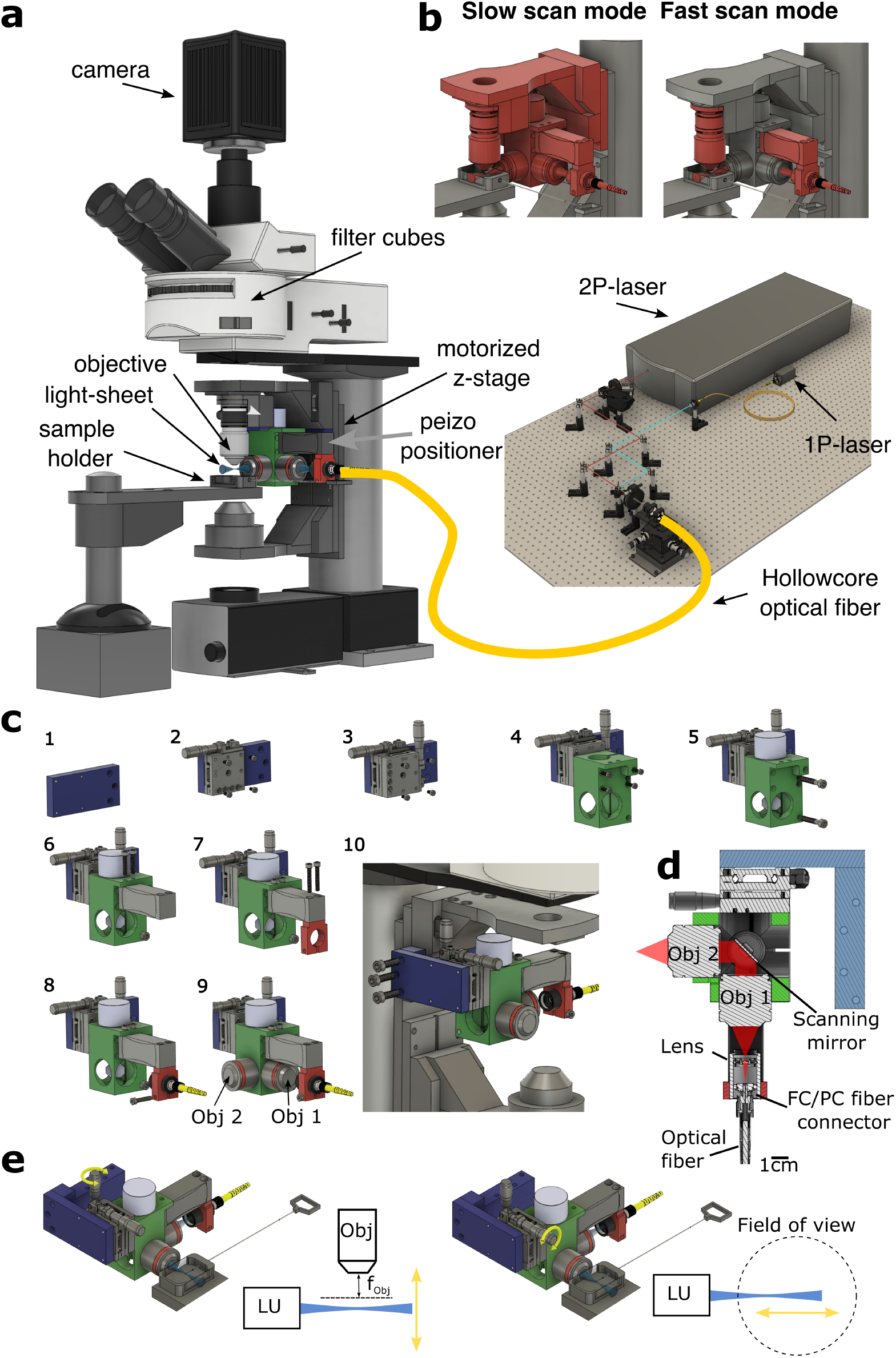
**(a)** 3D model of the complete setup. Explore the model interactively at https://a360.co/3C0TOAm and https://a360.co/434IdMt. **(b) Slow scan mode (Left):** The unit is mounted on a motorized stage that moves the objective, enabling slow z-scans through the sample while keeping the light sheet in the imaging focal plane. **Fast scan mode (Right):** Synchronized actuation of the piezo positioners, which hold the objective and fiber, enables fast, high-precision z-scans. **(c)** Step-by-step construction plan of the light-sheet unit. Additional construction plans with detailed explanations are available online at https://github.com/LJPZebra/OLU. **(d)** Cross-sectional view of the light-sheet unit, illustrating the galvanometric mirror that scans the laser beam in the sample plane to generate the digitally scanned light sheet. **(e)** Alignment procedure via mechanical manipulation of the entire light-sheet unit. **Left:** The unit (LU) is moved vertically using a mechanical z-stage (indicated by the yellow arrow) to align the light sheet with the imaging objective’s focal plane. **Right:** The LU is moved horizontally using a mechanical x-stage (indicated by the yellow arrow) to position the light-sheet waist at the center of the imaging objective’s field of view.

Our light-sheet unit (Figure 1c,d) comprises two objectives arranged in a one-to-one telescope configuration. The first objective collimates the laser beam from the fiber, and a galvanometric mirror at the back focal plane of the second objective pivots the beam, thereby sweeping it laterally to generate a digitally scanned light sheet. A key advantage of this light-sheet design is its minimal alignment requirement. During assembly, only two primary adjustments were needed: (1) coarse positioning of the galvanometric mirror at a 45° angle (with fine-tuning via the controller offset) to center the laser beam into the second objective (Obj 2), and (2) adjusting the distance between the fiber and the collimating objective (Obj 1) to ensure proper collimation. Once mounted on the microscope stand, no further internal alignment was required to position the light sheet within the focal plane of the imaging objective or to center its waist in the field of view. Due to the compact design, the final adjustments were easily made by translating the entire unit along the z-axis (to align the light sheet with the focal plane) and the x-axis (to center the waist) (see Figure 1e).

The unit is mounted via aluminum brackets to the microscope’s motorized vertical translation stage used for focusing (see Figure 1b, c). This configuration ensures that the detection objective and the illumination unit move synchronously during focus adjustments, simplifying volumetric scanning. Slow volumetric recordings are achievable via the built-in microscope software controlling the objective stage (see Video S2 and Figure 1b left). For fast volumetric scanning, two piezo actuators enable high-speed, synchronous motion of the imaging objective and the fiber outlet (see Video S3 and Figure 1b right).

A key advantage of our dual light-sheet system is the delivery of all laser sources through a single broadband optical fiber. This approach enables sharing a costly femtosecond laser (required for two-photon imaging) with other systems, thereby reducing the overall cost. Moreover, fiber delivery enhances flexibility and simplifies alignment. The entire unit can be manually translated along the z-axis to align the light sheet with the focal plane, and along the x-axis to center the light-sheet waist (see Videos S4 and S5, and Figure 1e). This alignment procedure is straightforward in one-photon mode, and no additional alignment is needed in two-photon mode because the fiber inherently coaligns the laser beams.

### B. Broadband Delivery of Both CW and Femtosecond Pulses via a Hollow-Core Negative Curvature Fiber

#### Characteristics of the Negative Curvature Hollow Core Fiber

We developed a negative curvature hollow-core (HC-NCF) that exhibits optical characteristics ideally suited for laser delivery in the context of one- and two-photon light-sheet microscopy (Figure 2). The fiber has an outer diameter of 200 ± 2 *µ*m and a fiber coating diameter of 600 ± 30 *µ*m. Its unique design features an air-filled 30 *µ*m-in-diameter core surrounded by a ring of eight non-contacting cladding tubes, each with an approximate diameter of 10 *µ*m. This hypocycloid negative curvature fiber structure enables the creation of a negative curvature core contour, which is essential for achieving the desired optical properties. Remarkably, this fiber exhibits two broad transmission bands that are well suited for one-, two-, and in principle also three-photon imaging (see Figure 2a). The first transmission band spans the visible spectrum from 400 to 535 nm (I), while the second transmission band covers the near-infrared range from 700 to 1500 nm (II). Within these spectral bands, the fiber demonstrates low loss, measuring less than 300 dB/km (equivalent to a transmission efficiency of over 93 % for a one-meter-long fiber), and minimal pulse dispersion, with less than 1 dB/(km·nm) of the spectral pulse width. Furthermore, this fiber is commercially available as patch-chord cable with standard FC/PC connectors, allowing for convenient disconnection and reconnection to the optical setup without the need for a subsequent realignment on either end. The small core diameter, in comparison to the large infrared wavelength, ensures near single-mode guidance while preserving the Gaussian beam properties of the laser at the fiber output. Our measurements indicate M^2^ values of 1.23 and 1.18 at laser wavelengths of 1030 nm and 515 nm, respectively. The near field and far field profiles exhibit high symmetry, with minimal ellipticity of 97.25 %, and 85.57 %, respectively (see Figure 2c). The mode field diameters at 1/*e*^2^ are 23 ±1 *µ*m and 26 ±1 *µ*m respectively, corresponding to a fiber numerical aperture of NA ≈ 0.02. In addition, the HC-NCF fibers demonstrate a high damage threshold due to negligible interactions between the light and the fiber material. In our tests, we observed no fiber damage even at peak powers of 40 *µ*J (1030 nm) and 10 *µ*J (515 nm) for 400 fs pulses, corresponding to average laser powers of 20 W and 4 W, respectively. These characteristics highlight the fiber’s ability to deliver high-quality pulsed Gaussian beams across a broad spectral range, ensuring high transmission efficiency, and low dispersion, and making it an excellent choice for high-resolution multiphoton microscopy applications.

**Fig. 2.**
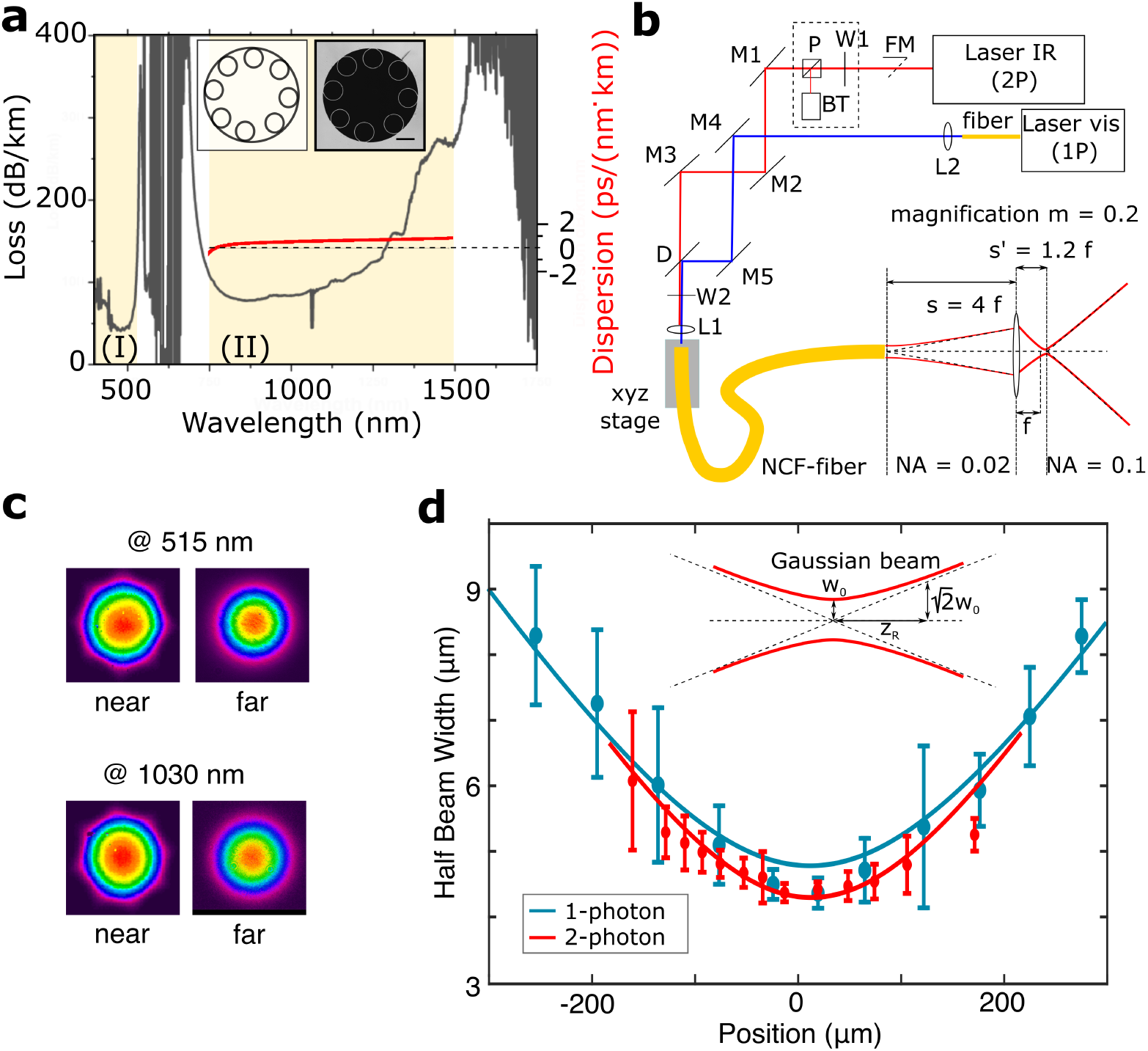
**(a)** Attenuation and dispersion spectrum of the hollow-core negative curvature fiber. Two transmission bands (loss *<* 300 dB/km) are highlighted in yellow. **Inset (left):** Schematic cross-section of the hypocycloid fiber with eight cladding tubes. **Inset (right):** Electron micrograph (EM) of the fiber cross-section (scale bar: 10 *µ*m). **(b)** Schematic of the optical path for coupling the laser source into the fiber and expanding the fiber output’s numerical aperture to match that of the collimation objective (Obj1 in Figure 1d). **(c)** Near-field and far-field beam profiles of the fiber output at 515 nm and 1030 nm wavelengths. **(d)** Measured axial light-sheet profile in the sample plane.

#### Fiber coupling: optical path and procedure

We achieved laser coupling into the fiber using a femtosecond pulsed Ti:Sapphire laser (MaiTai, Coherent, USA), along with three mirrors (M1-3), a dichroic (D), and a coupling lens (L1, f=40mm, Thorlabs, AC254-040-B-ML, see Figure 2b). The fiber was held in place by a differential xyz-translation stage. The continuous blue laser (488 nm, Oxxius, France) was coupled into the same hollow core fiber through the aforementioned dichroic using two additional mirrors (M4-5) and a collimation lens (L2). For efficient optical coupling and suppression of higher laser modes, the width of the laser focus projected onto the fiber input side must match the mode field diameter of the fiber. For the femtosecond laser, we positioned the coupling unit at a distance from the laser source where the slightly diverging laser beam reached the desired beam diameter, necessary to match the mode field diameter in combination with the chosen coupling lens. For the blue laser, we adjusted the beam diameter by selecting the focal distance of the collimation lens with the formula: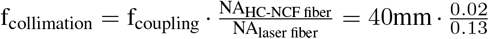, which boils down to the fact to choose the collimation lens focal distance such that the collimated laser beam diameter matches the diameter of the infrared beam that we previously coupled into the fiber using the chosen coupling lens. At a wavelength of 915 nm, corresponding to the central two-photon absorption peak of GFP and its calcium-sensitive derivative GCaMP, we delivered 100 fs laser pulses through 1,5 m fiber length with 98% power transmission efficiency and minimal pulse dispersion of 28 nm (1 dB/km·nm), which we fully precompensated with the Deepsee element of the MaiTai laser source. At 488 nm, the one-photon excitation maximum of GFP and GCaMP, we achieved a transmission efficiency of 75 %.

Two-photon light-sheet microscopy requires not only the delivery of high laser power through the fiber but also precise control of the laser’s polarization state. This is critical because fluorescence emission occurs predominantly perpendicular to the fluorophore’s dipole moment—and thus to the excitation light’s polarization plane (28). Consequently, if the laser is polarized perpendicular to the light-sheet plane, fluorescence is emitted mainly orthogonal to the detection objective’s optical axis, resulting in reduced collection efficiency (29). To optimize the fluorescence signal, we incorporated a half-wave plate before the fiber coupling unit to carefully adjust the laser polarization. This fine-tuning maximizes fluorescence emission and ensures proper alignment with the detection objective. It is important to note that once the optimal polarization is set, the fiber must remain free of bending or twisting. Cycloid-type negative curvature fibers are not polarization-maintaining (24), and mechanical deformation can alter the polarization state through rotation and changes in ellipticity.

#### Matching the NA of the light-sheet unit

Our compact light-sheet unit relies on an optical fiber to deliver the laser directly into the system. At the fiber outlet, the laser beam is first collimated by an objective and then focused onto the sample by a second objective (see Figure 1d). This configuration forms a one-to-one telescope that maintains unity magnification, projecting the fiber output directly onto the sample so that the light-sheet waist precisely matches the fiber’s mode field diameter.

In the one-photon implementation described by Migault et al., a single-mode optical fiber with a mode field diameter of 2.8–4.1,*µ*m at a 488,nm wavelength was used. This setup provided an ideal light-sheet waist and z-resolution, well-suited for cellular-resolution microscopy.

In contrast, our multiphoton implementation employs a hollow-core fiber with a considerably larger mode field diameter of 23,*µ*m—about five times larger than that of the single-mode fiber. This increase results in a reduced numerical aperture of 0.02. To recover the resolution of the one-photon configuration, we introduced an additional lens immediately after the fiber. This lens demagnifies the laser waist by a factor of five—reducing it to less than 5,*µ*m—while simultaneously increasing the beam divergence to match the collimation objective’s numerical aperture (NA 0.1) (see Figure 2b).

To achieve the targeted demagnification factor (m = 1/5 = 0.2), we positioned a lens with a focal length of f = 2,mm at a distance of *s* = *f* · (1*/m* −1) = 8 mm from the fiber outlet (see Figure 2b). This placement refocuses the laser to a new position at *s*′ = *f* · [1 + 1*/*(*s/f* −1)] = 2.4 mm beyond the lens, yielding a beam waist of 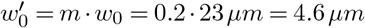. Additionally, the lens scales the laser divergence angle to m·0.02 = 0.1, which matches the numerical aperture of the collimation objective. Notably, the lens has a numerical aperture of 0.5, ensuring the laser beam is not clipped.

Finally, the compact lens can be directly affixed to the optical fiber outlet using a small lens tube. This add-on, termed the “NA-expander,” allows for seamless interchangeability with a standard single-mode fiber configuration.

### C. Imaging Performance

To demonstrate the system’s uncompromised resolution and speed, we recorded brain-wide 3D stacks of an immobilized larval zebrafish at cellular resolution using one- and two-photon imaging modes (Figure 3g and Video S6). We used transgenic zebrafish larvae expressing GCaMP6f under a pan-neuronal promoter. At six days post-fertilization, the larvae possess a largely transparent body, making them ideal for high-resolution microscopy. The specimens were embedded in low-melting-point agarose within a capillary and oriented so the brain faced the detection objective.

**Fig. 3.**
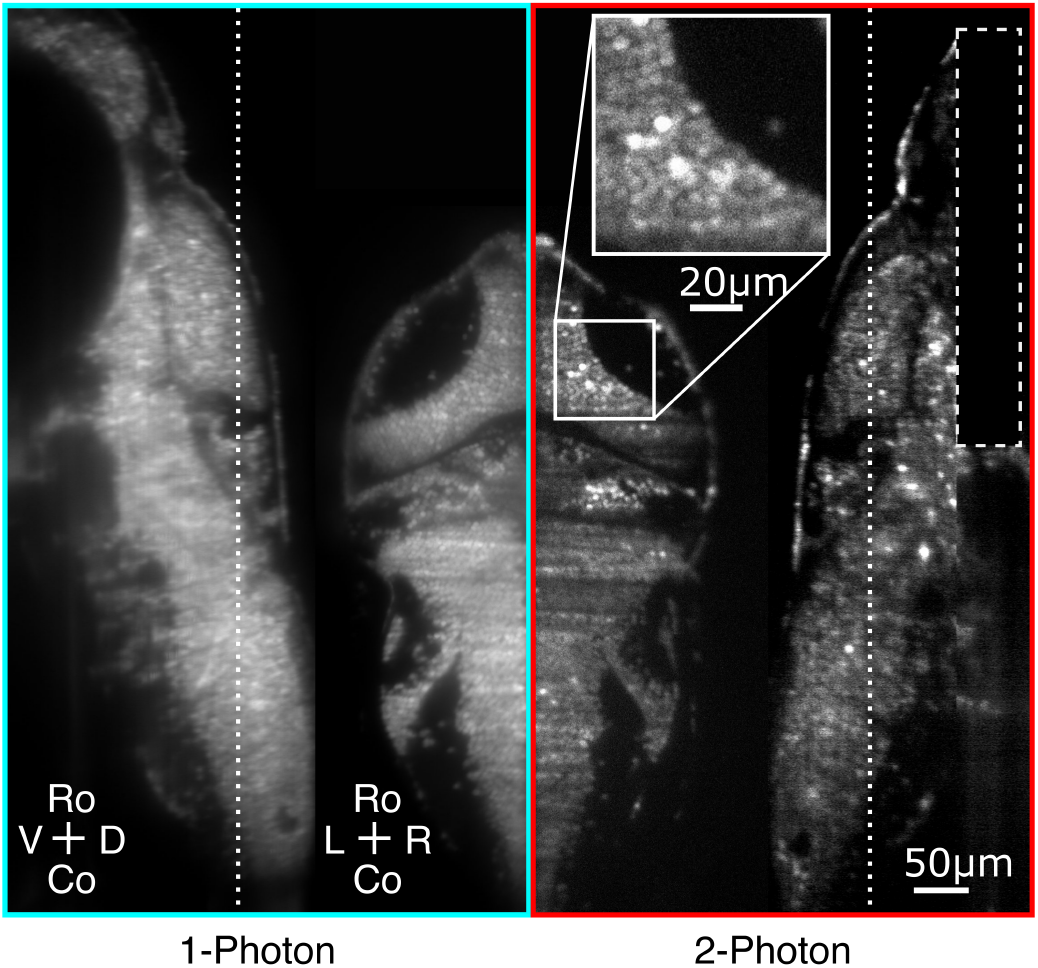
Volumetric imaging of a larval zebrafish brain in one- and two-photon modes. A 6 dpf zebrafish larva expressing pan-neuronal, nucleus-localized GCaMP6f (Elav3-H2B-GCaMP6f) was imaged. Both Sagittal and horizontal sections from the 3D volume are shown for each mode. In the sagittal section, the dashed line indicates the z-position of the horizontal plane shown. The inset highlights single cells resolved in the two-photon mode. In the two-photon mode, the light sheet was partially blocked during recording of ventral layers (dashed rectangle) to prevent it from striking the highly pigmented eyes, which would otherwise absorb excessive high-power infrared light, leading to heating and potential damage. The complete volumetric stack is available in Video S6. Abbreviations: Ro, rostral; Co, caudal; R, right; L, left.

For single-photon imaging, the sample was illuminated with a 488 nm continuous-wave laser light-sheet. In two-photon mode, we employed an ultrafast laser (100 fs pulses at 80 MHz) centered around 915 nm and relayed through the hollow-core fiber. Volumetric scans were achieved by synchronously moving the microscope objective and the light-sheet along the vertical axis using a fast, high-precision scan mode.

We acquired stacks from ventral to dorsal through the entire brain of a 6-day-old larva with 1-µm z-increments (see Video S6 and Figure 3). Both imaging modes resolved single neurons across the entire brain. Although single-photon imaging produced relatively bright fluorescence signals, it exhibited slightly increased background at deeper layers. In contrast, two-photon imaging offered superior optical sectioning and reduced out-of-focus fluorescence, particularly in highly scattering regions—consistent with previous observations that infrared excitation penetrates deeper into larval zebrafish brains with minimal scattering.

Furthermore, we conducted rapid whole-brain functional imaging of spontaneous neural activity (see Video S7). This approach enabled us to monitor real-time neural dynamics and capture transient events with high temporal fidelity across the entire brain. Delivering several hundred milliwatts of laser power into the sample—a requirement for fast functional two-photon light-sheet imaging—was made possible by the fiber’s high transmission efficiency and damage threshold, further underscoring the versatility of our system. Overall, these findings demonstrate that our compact one- and two-photon light-sheet unit, based on laser delivery via a negative curvature hollowcore fiber, excels in both high-resolution structural imaging and rapid functional imaging. In conclusion, our system offers an integrated solution that combines precision and speed, making it a powerful tool for advanced neurobiological research.

## Discussion

Our results demonstrate that integrating a compact, fiber-coupled light-sheet unit, which employs a negative curvature hollow-core fiber, can transform a standard upright commercial microscope into a dual-mode one- and two-photon LSFM system. This approach overcomes several technical and financial barriers commonly associated with multiphoton microscopy.

First, the system enables sharing a single femtosecond laser source across multiple setups through hollow-core fiber patch cables. Laboratories already equipped with a pulsed ultrafast near-infrared laser for two-photon imaging can extend multiphoton capabilities to an additional microscope without investing in a second expensive laser. Second, fiber-based laser delivery greatly simplifies optical alignment. Because the beam exits the fiber pre-aligned, the unit can be easily translated and rotated to align the light sheet relative to the imaging focal plane without the need to adjust free-space optical elements. Moreover, when incorporating a second laser line, the user only needs to optimize the input coupling. Once initially set up, the system eliminates the need for repeated free-space alignments, reducing maintenance effort.

The broadband transmission of the negative curvature fiber further enhances the system’s flexibility. It supports wavelengths spanning from the visible to the near-infrared spectrum (e.g., 700–1050 nm), enabling the excitation of diverse fluorophores for one- and two-photon imaging, and even facilitating three-photon excitation for deeper tissue penetration when the optical path is appropriately optimized. The same fiber can transmit visible light for single-photon fluorescent excitation, and tunable sources—such as optical parametric oscillators—could be integrated to expand the range of compatible fluorophores and experimental designs. This versatility makes the unit well-suited for advanced applications, including multispectral functional imaging and combined optogenetic photostimulation, where adapting the light sheet wavelength can help reduce cross-talk with the stimulation. In addition, the fiber’s optical properties and broadband transmission spectrum render it ideal for implementation in other microscopy modalities.

Our demonstration of brain-wide calcium imaging in larval zebrafish underscores the system’s capability for rapid, volumetric functional imaging. The compact design and fiber coupling do not compromise microscope resolution or speed. While our validation was performed on zebrafish larvae, the system is readily adaptable to other small model organisms, embryonic samples, organoids, or ex vivo tissues with comparable optical resolution.

Beyond hardware performance, the open-source nature of our design promotes accessibility and further innovation. We provide detailed assembly instructions, mechanical drawings, 3D models, and software for coordinating the galvanometric mirror, objective motion, and camera acquisition. Importantly, by mounting the unit onto the vertical microscope translation stage, the light-sheet unit moves synchronously with the objective, enabling volumetric imaging using standard microscope control software or manual focusing without additional interfaces.

Moreover, the system is compatible with future enhancements. For instance, the fiber’s broadband transmission supports multicolor two-photon imaging (30); the digitally scanned light sheet can be paired with line-confocal detection to improve signal-to-noise ratios (31); and the fiber’s high damage threshold permits the delivery of high-energy pulses at low repetition rates, thereby increasing imaging speed while reducing phototoxicity (12).

In summary, our work provides a unified and flexible framework that brings two-photon light-sheet microscopy closer to a plug-and-play experience. By leveraging a single-fiber approach, we significantly reduce both the cost and complexity of building a two-photon LSFM setup, opening the door to broader adoption of multiphoton microscopy—even in laboratories traditionally limited to single-photon modalities.

## Methods

### Experimental model and subject details

All experiments were conducted on zebrafish nacre mutants aged 6-7 days post-fertilization (dpf). Larvae were reared in Petri dishes filled with E3 solution under a 14/10 hr light/dark cycle at 28 ^°^C. Fish were provided with powdered nursery food daily from 6 dpf. Calcium imaging experiments were performed in nacre mutant larvae expressing the calcium indicator GCaMP6f. The nearly pan-neuronal promoter elavl3 controlled the expression, and the H2B domain the localization of the sensor in the nucleus Tg(elavl3:H2B-GCaMP6f). The GCaMP6f line was provided by Misha Ahrens and published in Quirin et al. (32). The experimental protocols were approved by Le Comitée d’Ethique pour l’Expérimentation Animale Charles Darwin C2EA-05 (02601.01 and 32423-202107121527185 v3).

### Fiber Coupling Procedure

Coupling a laser into our hollow-core fiber is more challenging than coupling into standard fibers due to its low numerical aperture (NA_fiber_ = 0.02). To simplify this process, we developed a pre-alignment protocol (detailed instructions are available online at https://github.com/LJPZebra/OLU). Initially, a fiber-coupled visible laser was injected into the hollow-core fiber from the opposite end by connecting it via a fiber-to-fiber connector (Thorlabs, ADAF1) to our FC/PC-connectorized hollow-core fiber. In this configuration, the visible laser traversed the coupling optics in reverse order. The optical components were then pre-aligned so that the visible laser beam was collimated by the coupling lens and centered on the output aperture. Following this initial alignment, the visible laser fiber was disconnected from the hollow-core fiber outlet, a power meter was positioned at the outlet, and the femtosecond laser was activated at low power to prevent fiber damage during subsequent adjustments. Once the pre-alignment yielded sufficient laser transmission for reliable infrared measurements, the coupling was further fine-tuned until a transmission efficiency greater than 90% was achieved.

#### Characterization of the light-sheet profile

We followed the protocol presented in Wolf et al. 2015 (6) to characterize the light-sheet profile. In brief, we acquired a high-resolution stack of a sample containing sub-diffraction-sized fluorescent beads (FluoSpheres, 505/515, Molecular Probes, USA) with a diameter of 100 nm embedded in 2 % low melting point agarose. The z-spacing between consecutively imaged sections was set at 0.5 *µ*m. For each bead, the fluorescence intensity profile across the light sheet was fitted with a Gaussian function, from which we extracted the half width at 1/e^2^. This width, which depends on the bead’s position relative to the light-sheet waist, corresponds to the beam profile of the Gaussian beam generating the light sheet. To determine the light-sheet waist, *ω*_0_, we fitted the measured bead profile widths as a function of the position *x* along the light sheet using the Gaussian beam profile equations (Eq. 1 for the one-photon profile and Eq. 2 for the two-photon profile) as described in (6).

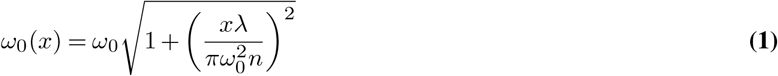

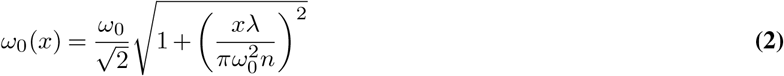

with the laser wavelength λ, the refractive index *n*, and the waist of the Gaussian beam *ω*_0_.

### Acquisition and Processing of Functional Light-Sheet Data

Image acquisition, mirror scanning, and objective translation were controlled and synchronized using National Instruments cards (NI 9402 and NI-9263, National Instruments) together with a custom MATLAB program (The MathWorks), which is available open source at https://github.com/LaboJeanPerrin/Lightsheet. Offline image pre-processing and extraction of calcium transients (.6.F/F) were performed in MATLAB following the workflow described in (9, 27). The acquisition parameters for the brain recordings shown in Figure 3g were as follows: For the one-photon recording, the interlayer interval was 1 *µ*m, the exposure time was 20 ms, the continuous laser power was approximately 1 mW, and the excitation wavelength was 488 nm. For the two-photon recording, the interlayer interval was also 1 *µ*m, the exposure time was 11 ms, the mean laser power in the sample was 710 mW, the pulse duration was 100 fs, the pulse repetition rate was 80 MHz, and the excitation wavelength was 915 nm.

#### Parts List

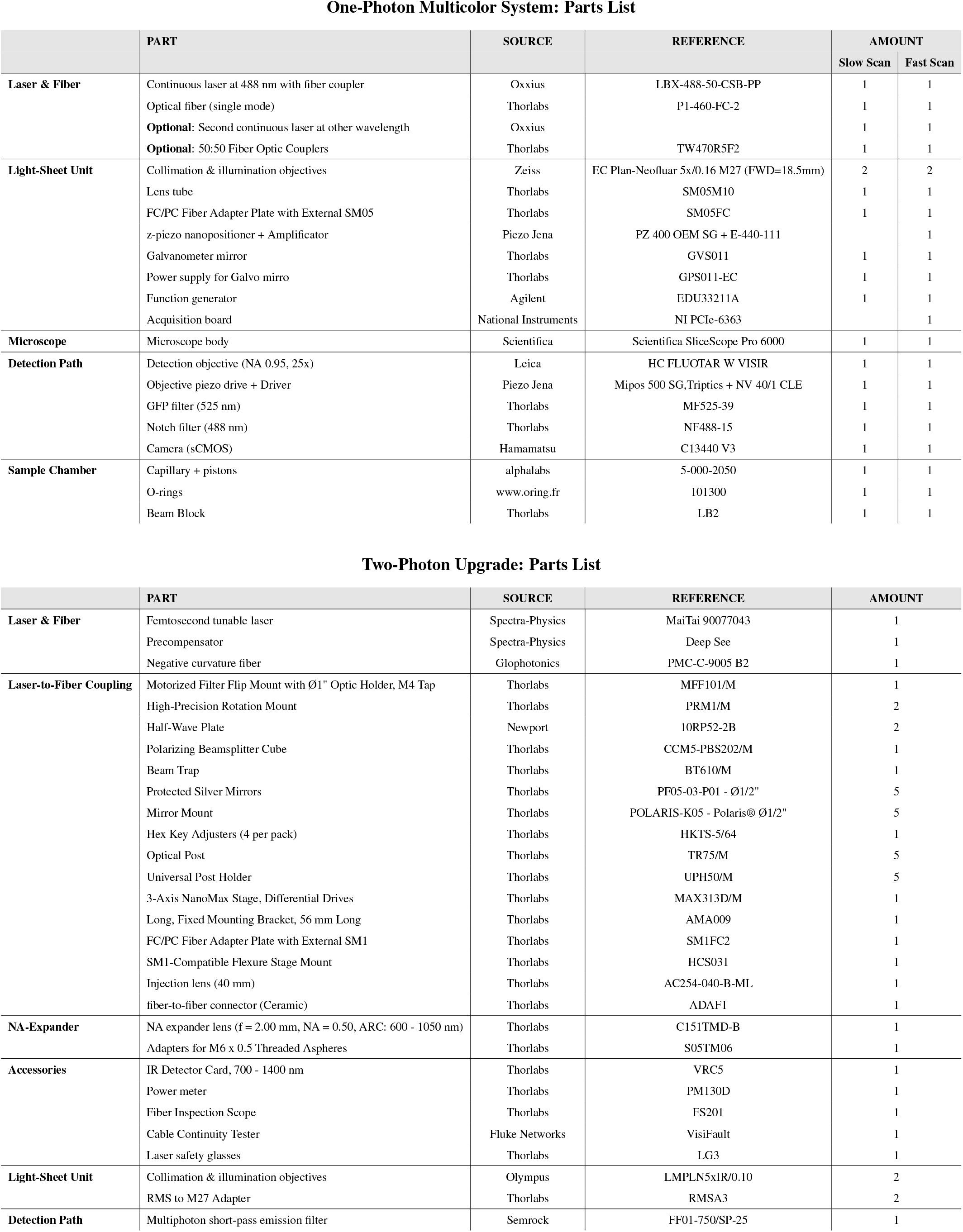

## ACKNOWLEDGEMENTS

The authors thank the IBPS fish facility staff for maintaining the fish. We thank Misha Ahrens for providing transgenic fish lines. We are grateful to Carounagarane Dore for his contribution to the design of the experimental setup. We thank Loic Royer for his valuable feedback and discussions related to the mansucript and the online material. This project has received funding from the European Research Council (ERC) under the European Union’s Horizon 2020 research innovation program, grant agreement number 715980. Furthermore, it was supported by grants from Région Ile-de-France, specifically by DIM cerveau et pensée and DIM-ELICIT. Moreover, the project received partial funding from the CNRS and Sorbonne Université. GM had a Ph.D. fellowship from the Doctoral School in Physics, Ile de France (EDPIF), HT had a Ph.D. fellowship from Ile de France (ARDoC 17012950).

## AUTHOR CONTRIBUTIONS

AH, TP, GM, HT, GD, and VB designed the project, performed research, analyzed data, and wrote the manuscript. B.B. developed and characterized the broadband optical fiber

## DATA AVAILABILITY

All data generated or analyzed during this study are included in this published article, its supplementary information files, and the online resources (https://github.com/LJPZebra/OLU).

## SUPPLEMENTAL INFORMATION

### Supplemental videos

**Video S1 Animated 3D model of the design of the light-sheet microscope**. The video sequence begins with an overview of the system, highlighting the sample positioning. It then demonstrates the manual focusing that automatically adjusts the light-sheet position to the sample plane. The video then zooms in on the light-sheet unit to showcase its precise mechanical adjustments, which optimize the light-sheet waist and improve image quality. Finally, the animation illustrates both the slow and fast scan modes.

**Video S2 Slow scan mode, manual or computer controlled**. This video demonstrates the slow volumetric scanning mode of the light-sheet microscope. The highlighted parts, shown in red, move together as a unit to perform the scan. These components are attached to the motorized stage of the microscope stand, which is typically used for controlling the focus of the microscope objective. The sample can be scanned via manual or computer-controlled stage adjustments while the light-sheet consistently illuminates the focal plane.

**Video S3 Fast High-Precision Volumetric Scanning Mode**. This video showcases the light-sheet microscope’s fast and high-precision volumetric scanning mode. The objective and optical fiber, highlighted in red, move together and are micro-positioned using piezo crystals, also depicted in red. These components are synchronized in their movement and controlled by a computer to perform a scan through the sample while continuously illuminating the focal plane with the light sheet.

**Video S4 Alignment of the light-sheet into the detection objective focal plane**. This video demonstrates the alignment process to position the light sheet precisely within the detection objective’s focal plane. The animation highlights the movement of the light-sheet unit along the z-direction, which can be adjusted using the z-stage control screw. An inset video on the right shows the focus changes as alignment is adjusted, with the laser visualized using a fluorescein solution. An inset on the left illustrates the impact of alignment on recording quality using a brain sample from a 6-day-old zebrafish.

**Video S5 Alignment of the laser waist within the field-of-view center**. This video demonstrates the alignment procedure for positioning the laser waist precisely at the center of the field of view. The animation highlights the movement of the light-sheet unit along the laser axis, which can be adjusted using the x-stage control screw. An inset video on the right shows the focus changes as alignment is adjusted, with the laser visualized using a fluorescein solution. An inset on the left illustrates the impact of alignment on recording quality using a brain sample from a 6-day-old zebrafish.

**Video S6 Example Anatomical Brain Recording in One- and Two-Photon Mode**. This video demonstrates dorsal-to-ventral scans through the brain of a 6-day-old larval zebrafish expressing the pan-neuronal calcium sensor GCaMP6f (Tg(Elav3-GCaMP6f)). The left panel shows a recording obtained in one-photon mode using a 488 nm laser, while the right panel displays a recording in two-photon mode at 915 nm. Both recordings were performed consecutively on the same fish to allow a direct comparison between the two imaging modalities. In the two-photon mode, the light sheet was partially blocked during recording of ventral layers to prevent it from striking the highly pigmented eyes, which would otherwise absorb excessive high-power infrared light, leading to heating and potential damage.

**Video S7 Spontaneous Brain Activity Recorded in Two-Photon Mode**. This video showcases the high-precision fast scan mode and its application in recording spontaneous brain activity in a 6-day-old larval zebrafish brain expressing the panneuronal fluorescence calcium sensor GCaMP6f (Tg(Elav3-GCaMP6f)). The video was recorded in two-photon mode at a wavelength of 915 nm. The video displays four brain slices simultaneously recorded at different heights, starting from the most dorsal layer, with the signals displayed represented as DFF. The video is shown at 10x speed for enhanced visualization.

### Printed version of the open-source online tutorial

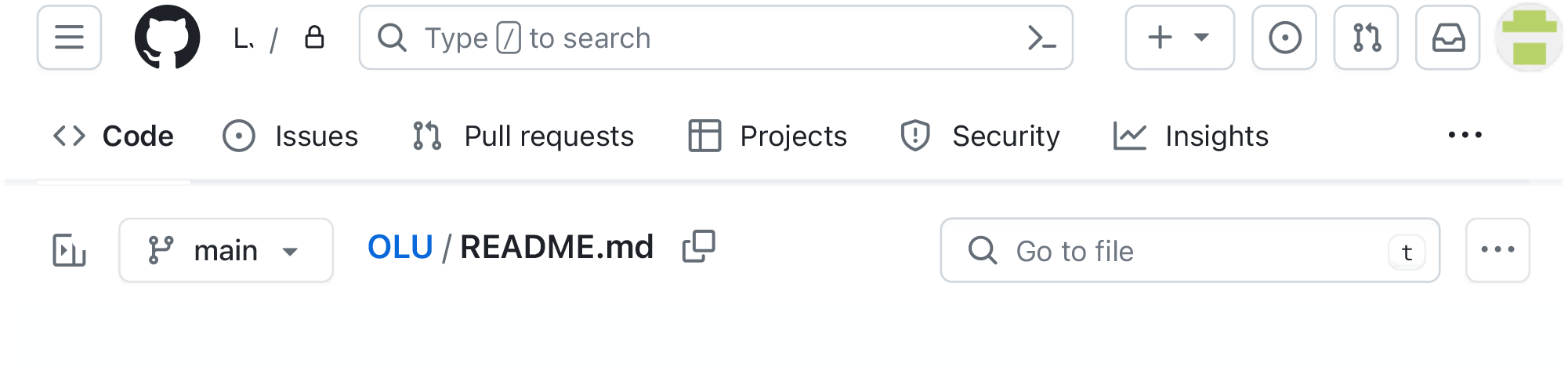

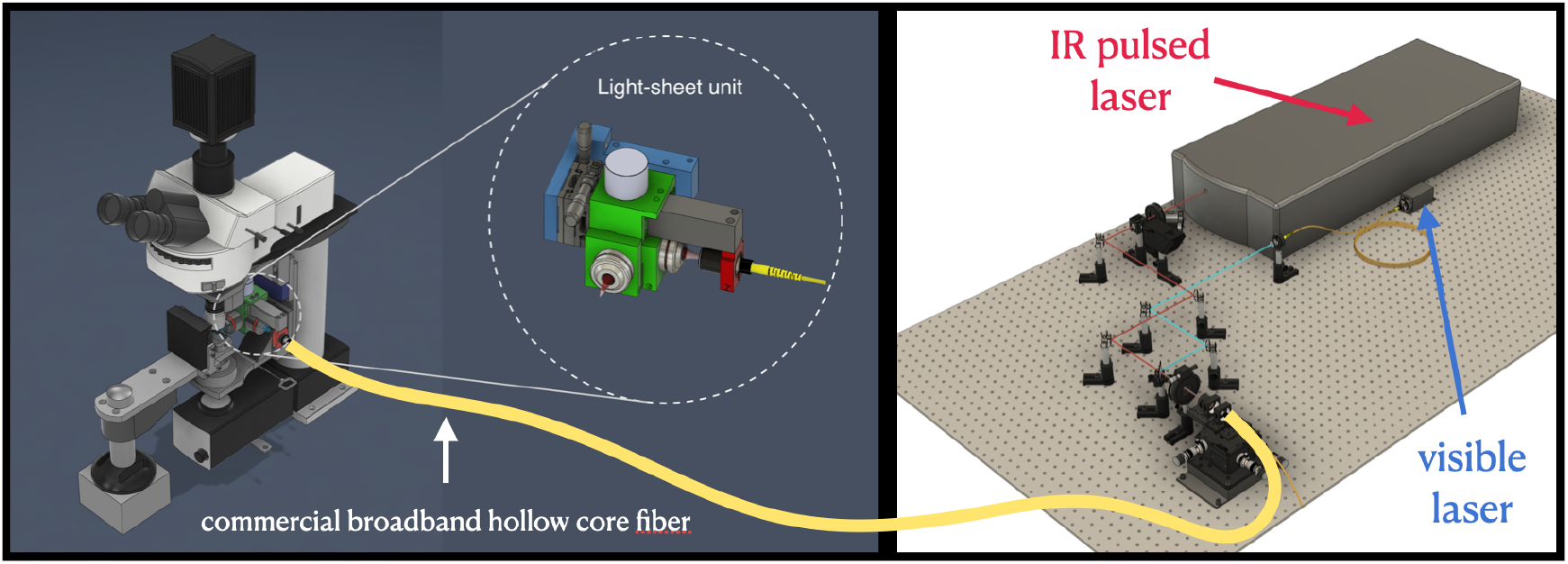

Two-photon light sheet microscopy offers great potential for a range of biological applications, but its practical implementation is impeded by the high cost of laser sources, the complexity of construction, and the challenges associated with adapting to existing microscope setups. Here, we release an open-source design that addresses these limitations by providing detailed building instructions for the transformation of a brightfield microscope into a versatile one- and two-photon light sheet system. Our design incorporates a specially designed broadband hollow core fiber, enabling the simultaneous utilization of an expansive pulsed laser source from another setup alongside a visible laser. This integration allows for uncompromised image resolution and speed. Furthermore, the design reduces the complexity of construction, alignment, and overall cost, thereby significantly enhancing the accessibility of this technology.

## BUILDING INSTRUCTIONS

Here we provide a comprehensive online tutorial that features step-by-step building instructions along with 3D animations, illustrations of the construction process, detailed information on the underlying physical principles, and rationale for components selection. In conjunction with the availability of commercially accessible and connectorized broadband optical fiber, our aim is to promote greater accessibility and affordability in the field of multiphoton light-sheet microscopy. To access this online resource, please follow the links provided below:

1. Build first a multicolor one-photon version of the microscope with fiber delivery of lasers in the visible spectrum with a standard single-mode optical fiber technology.
2. Upgrade the one-photon unit into a two-photon system by exploiting advanced hollow core fiber technology.

## Citation

If you use the hardware designs, please cite us:

A Versatile and Open Source One- and Two-Photon Light-Sheet Microscope Design

Thomas Panier*, Geoffrey Migault*, Antoine Hubert, Hugo Trentesaux, Benoît Beaudou, Georges Debrégeas, Volker Bormuth

## Explorable 3D model of the full system

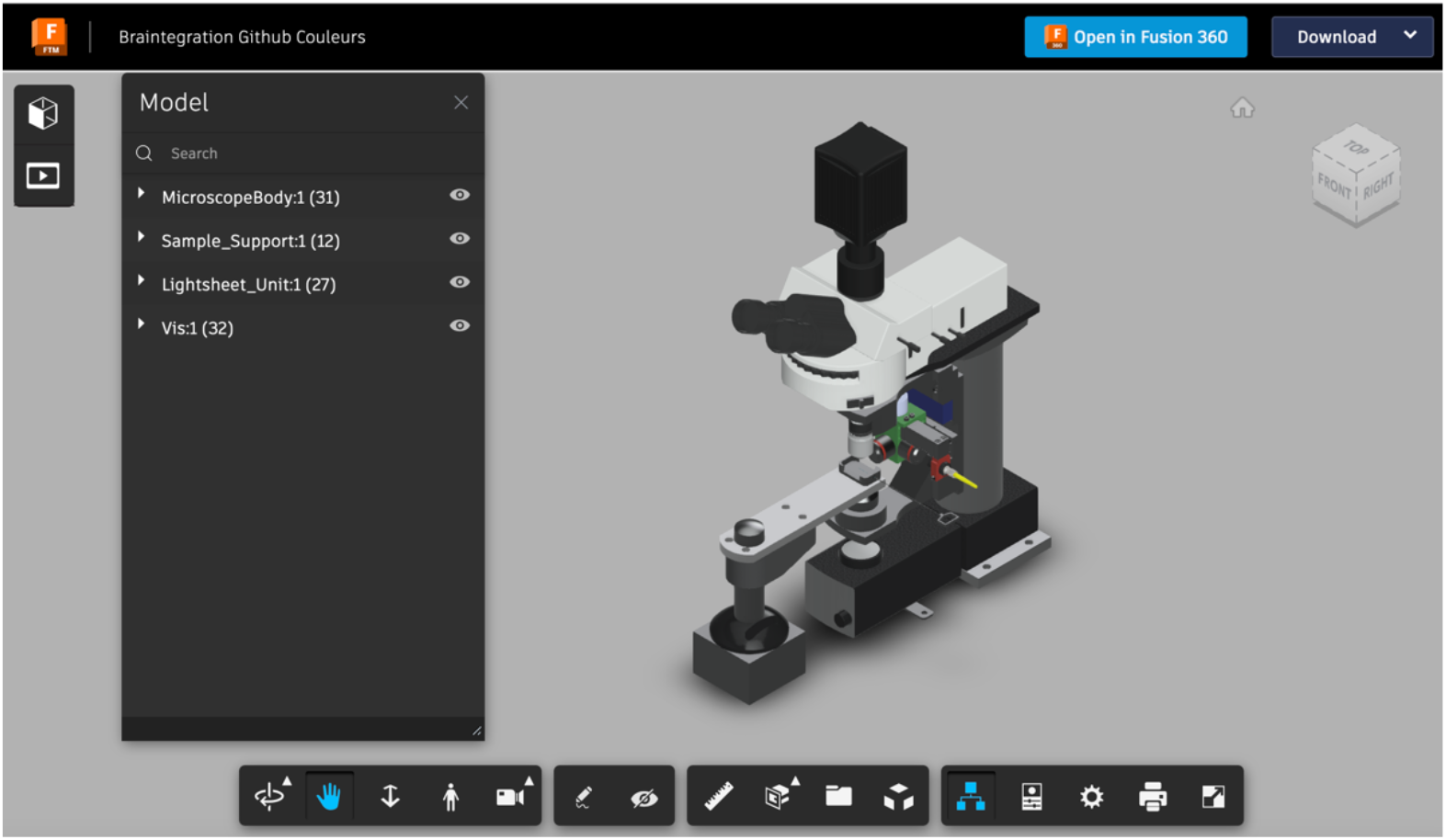

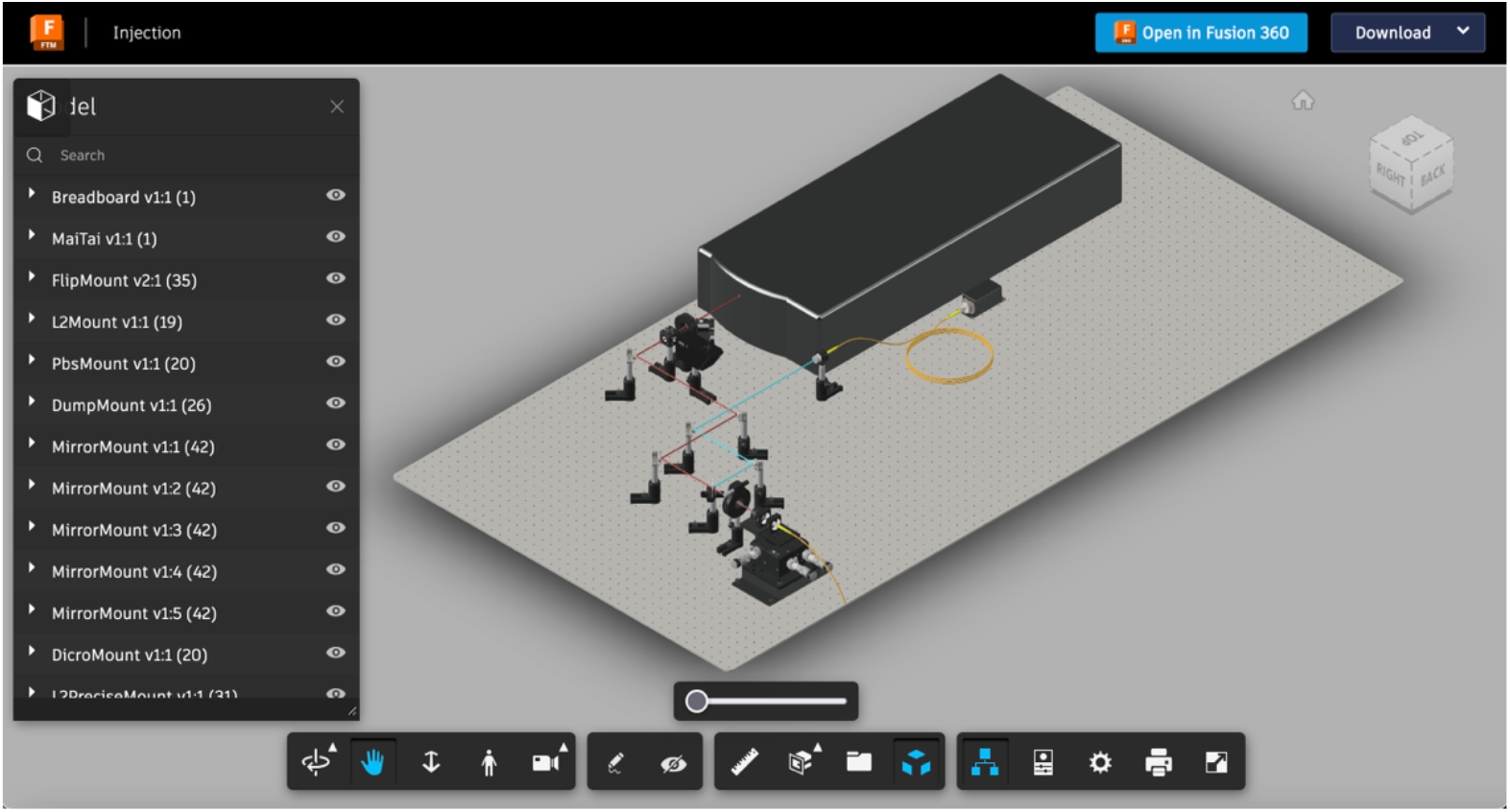

Click the images to open the model browser.

## 3D Animation of the functionality of the system

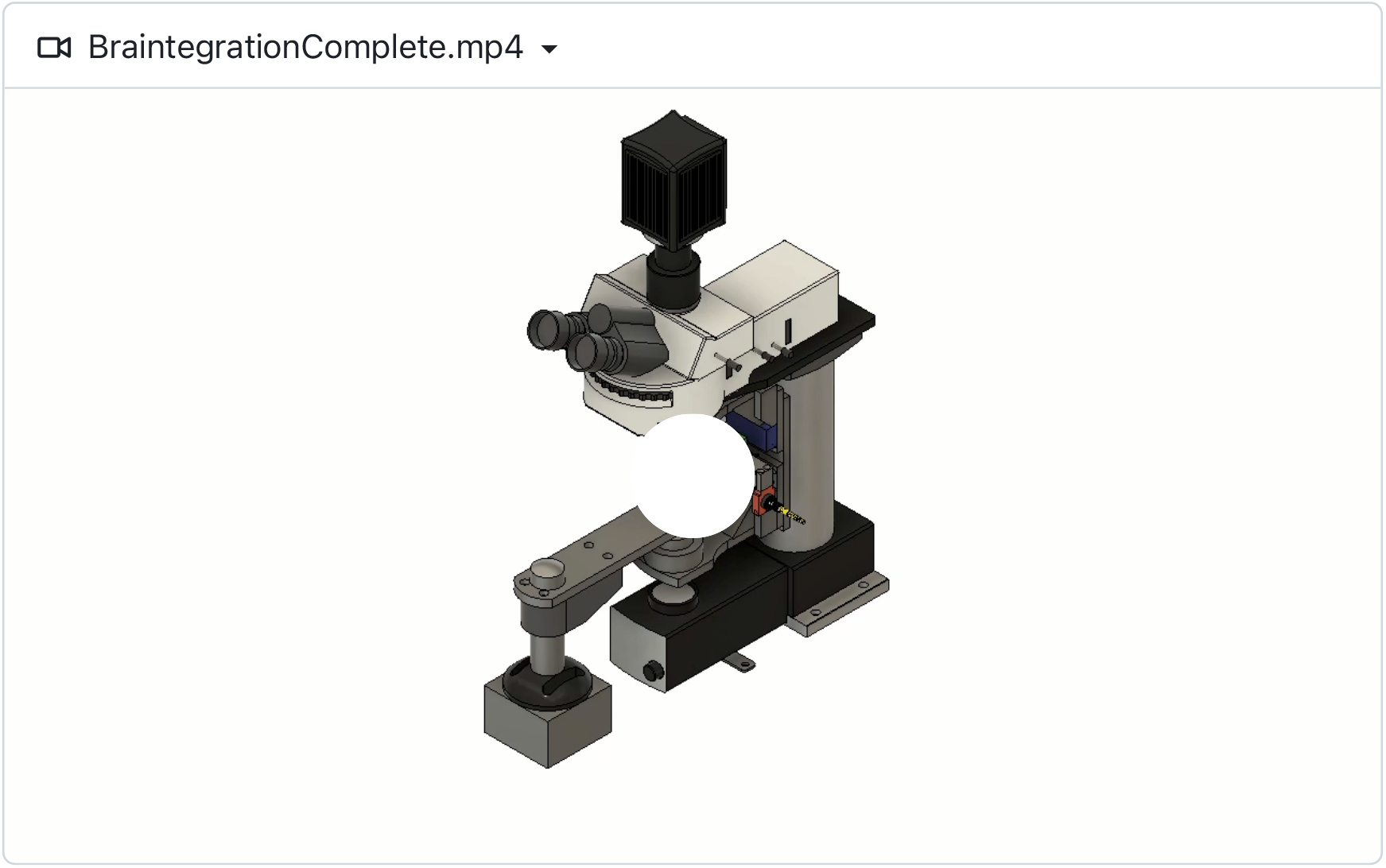

## Slow scan mode

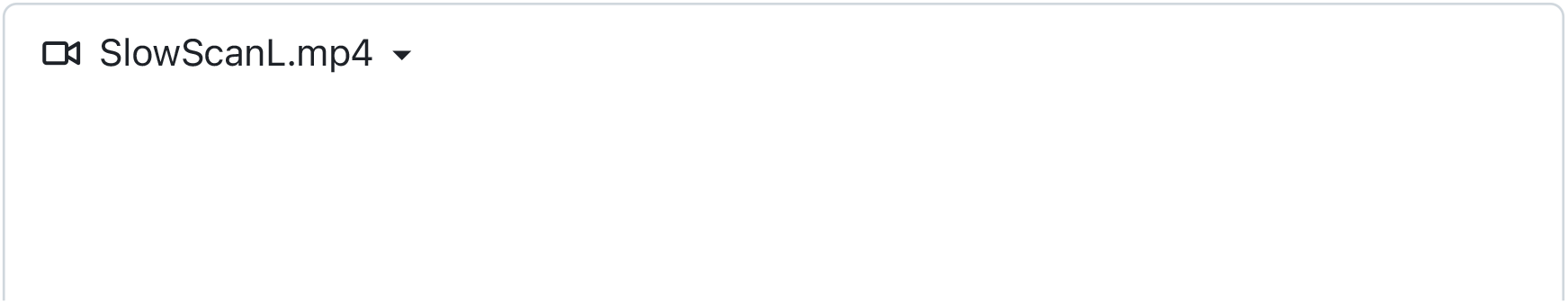
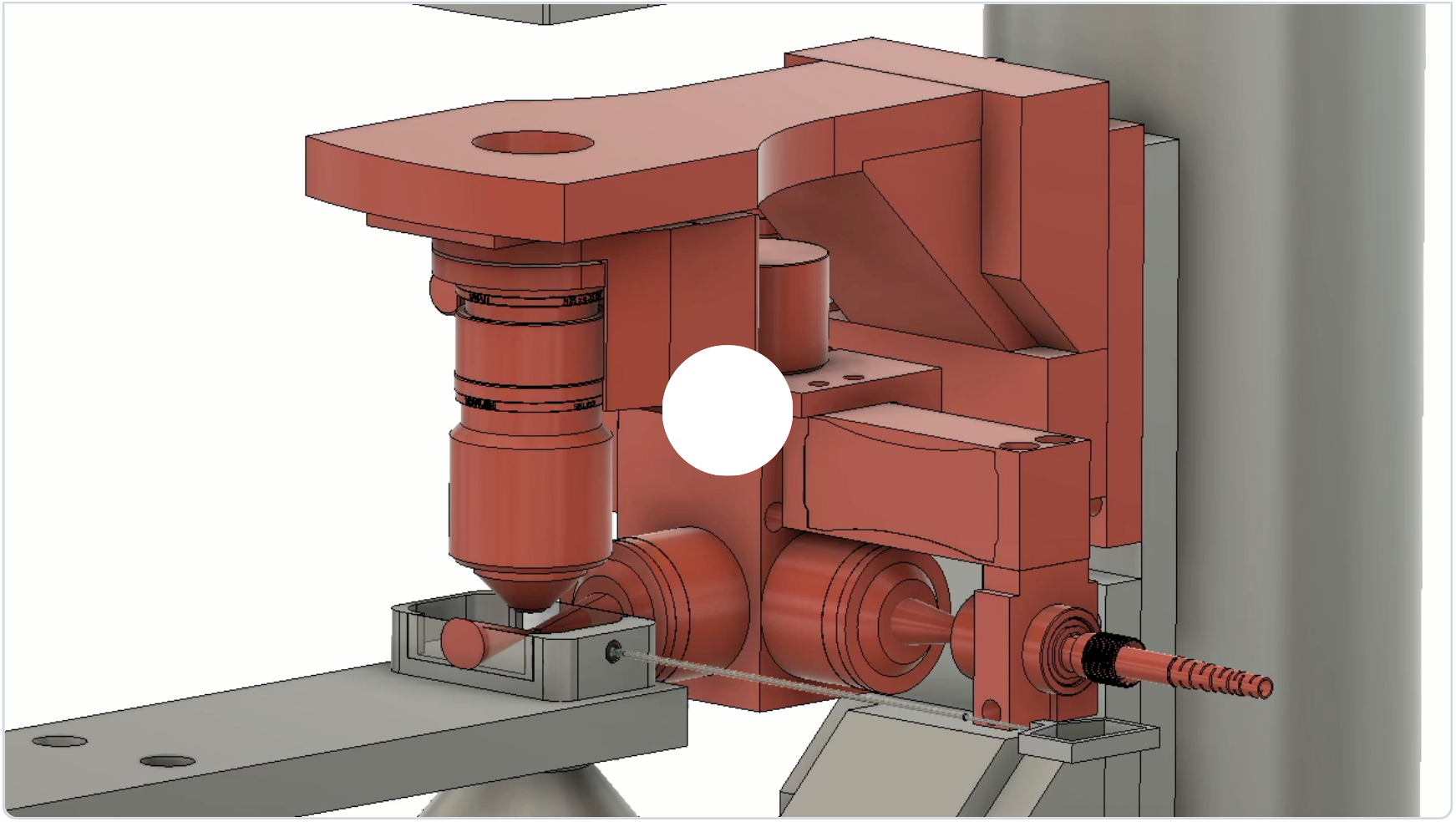

## Fast scan mode

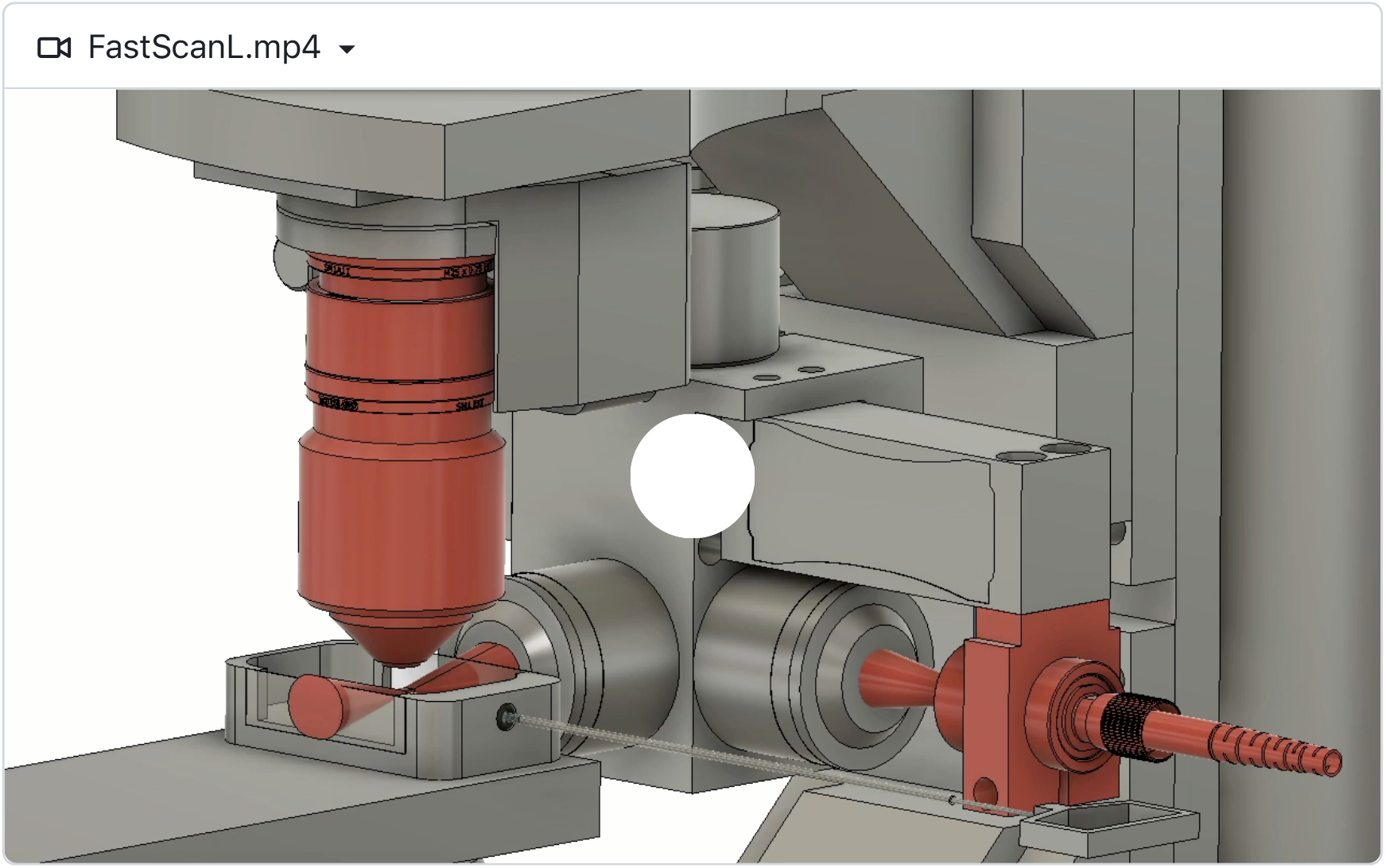

## High-resolution zebrafish brain scans (elav3:H2B-GCaMP6)

- Left: one-photon mode excited @ 488nm
- Right: two-photon mode excited @ 915nm

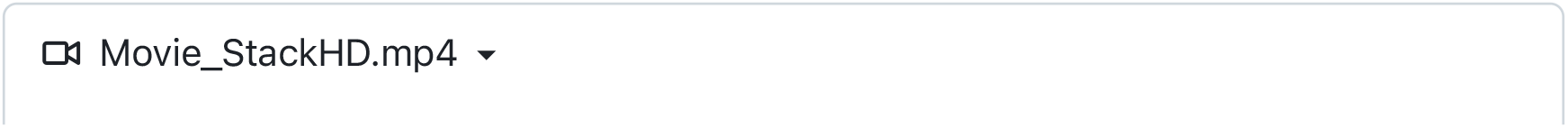
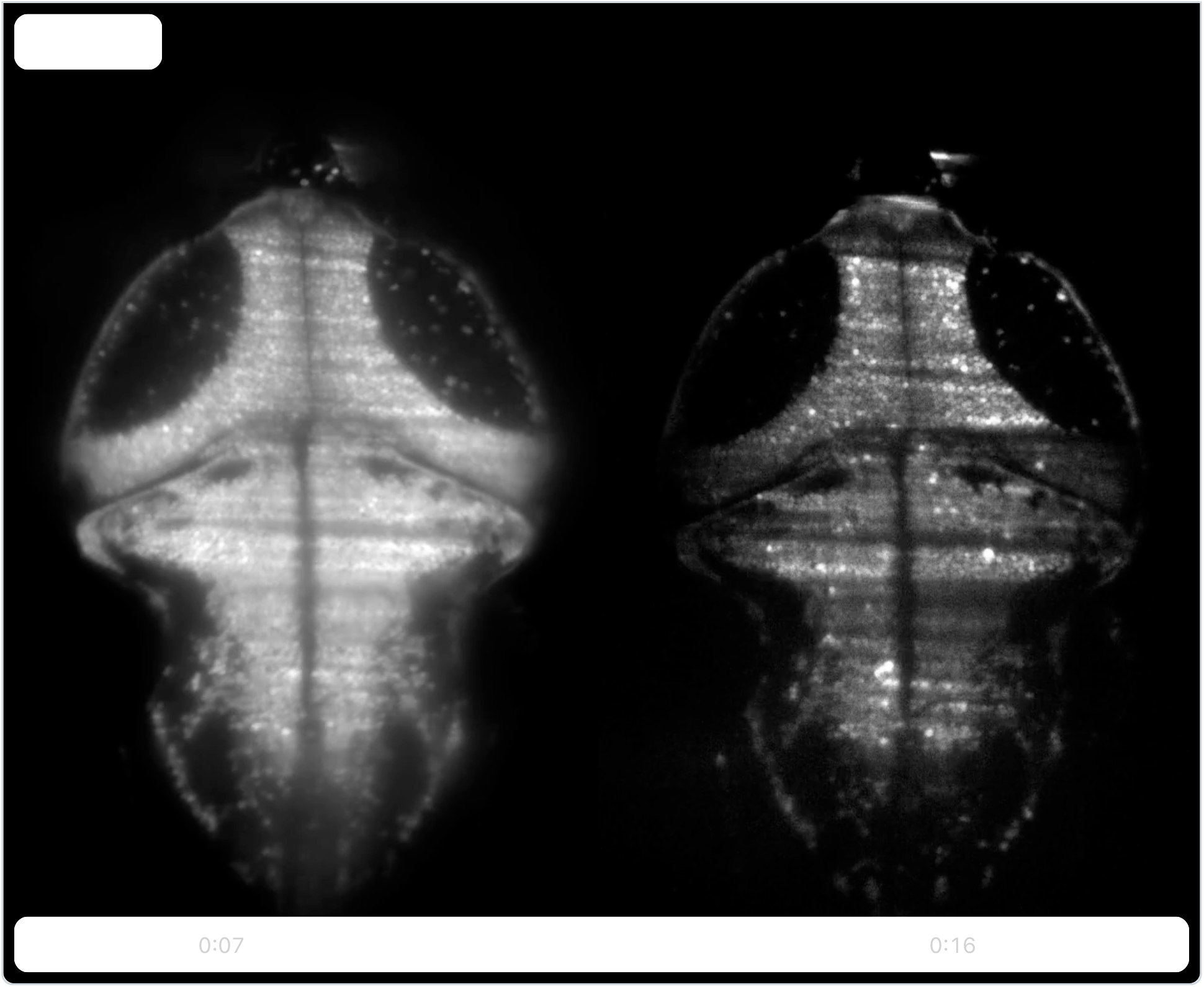

## Spontaneous brain activity recorded in a two-photon mode

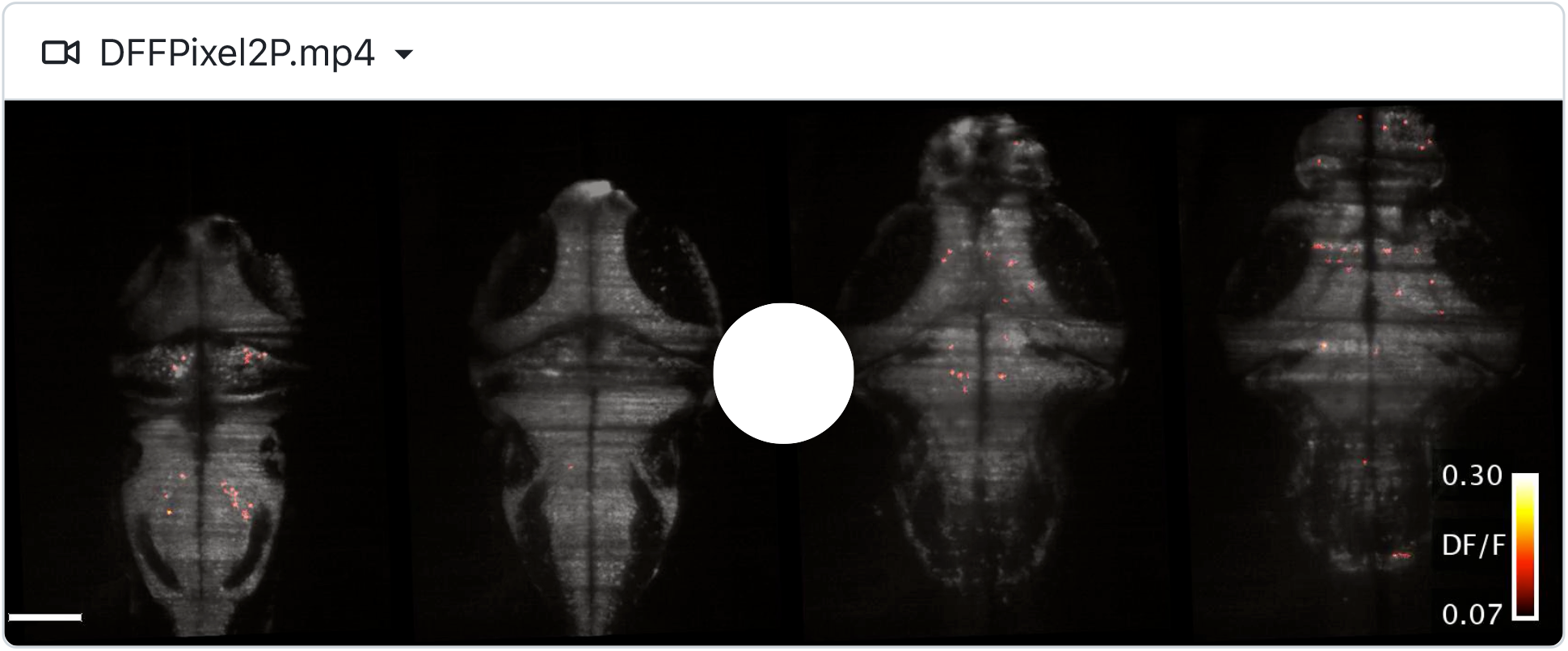

## CAD models

### CAD models

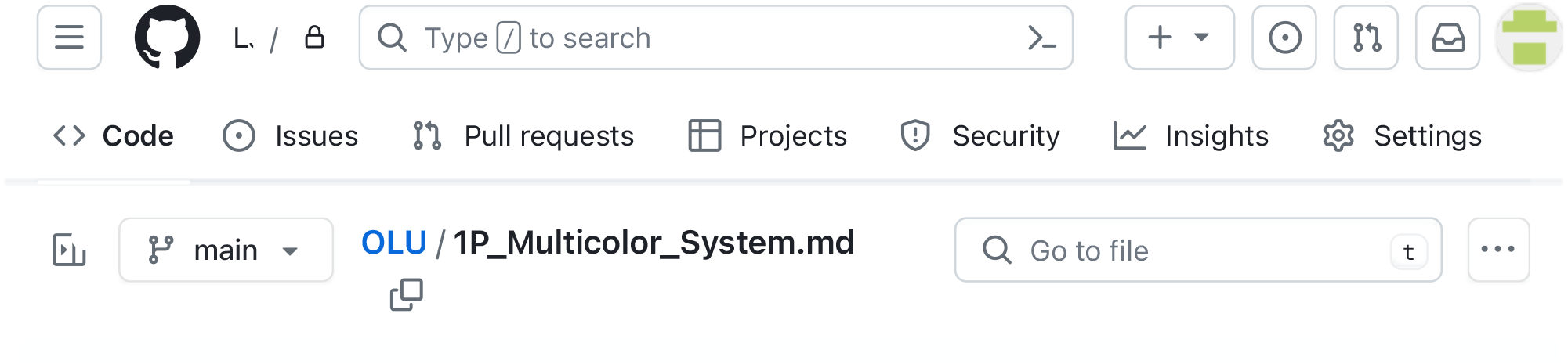

## The Multicolor One-Photon Light Sheet Design

This comprehensive guide provides detailed instructions for building the one-photon light-sheet system. Whether your ultimate goal is to construct the two-photon system or utilize the one-photon configuration, we highly recommend starting with the installation of the one-photon system. This sequential approach allows you to gain a solid foundation and a better understanding of the system’s components and functionality, which will facilitate a smoother transition to the advanced two-photon configuration if desired.

The construction of the system can be done on two levels. At the first level, the light-sheet unit is a standalone light source. This basic configuration allows you to perform light-sheet imaging experiments in the same manner as you would normally acquire epifluorescence images or stacks. At the second level, the concerted motion of the light-sheet and imaging objective is computer-controled and allows you to perform fast, high-precision volumetric scans. This advanced level of control opens up additional capabilities and enhances the versatility of the light-sheet unit for various imaging applications.

If your ultimate goal is to set up the two-photon version, you have the option to directly use optics optimized for infrared transmission in the initial one-photon installation. This strategic choice ensures compatibility and paves the way for a seamless transition to the two-photon configuration at minimal cost by avoiding the need for component replacements when upgrading to the two-photon version.

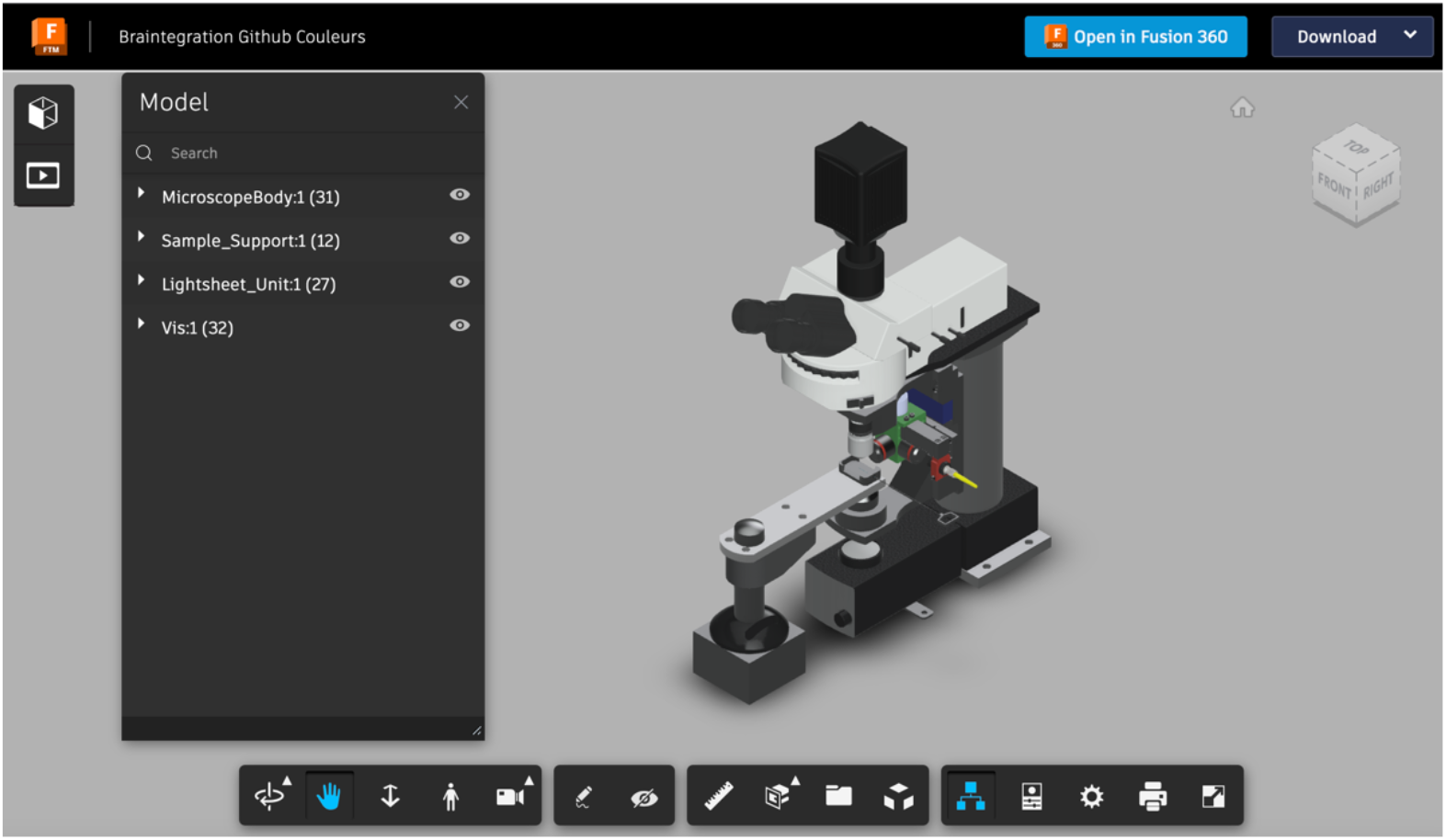

Click the images to open the model browser.

## Building Instructions

### Purchase List

List of parts

### Custom Parts

#### Parts to Send for Milling

These parts are milled from an aluminum block. If you do not have a mechanical workshop in-house then you can send the *.step files that we provide below to an online milling service. For parts with threaded holes, join the mechanical drawings to the .step file.

- The light-sheet unit central cube: View the 3D model or download the CAD model as a step file and mechanical drawings.
- The fiber holder : View the 3D model or download the CAD model as a step file and mechanical drawings.
- Adaptor plate 1 : View the 3D model or download the CAD model as a step file and mechanical drawings.
- Adaptor plate 2 : View the 3D model or download the CAD model as a step file and mechanical drawings.
- Custom bracket : View the 3D model or download the CAD model as a step file and mechanical drawings.
- Sample chamber holder : View the 3D model or download the CAD model as step file and mechanical drawings.
- T-bracket to hold the beam trap : View the 3D model or download the CAD model as step file and mechanical drawings.
- Piezoactuator dummy : View the 3D model or download the CAD model as step file and mechanical drawings. This piece is only necessary if you do not want to install the piezo actuator required for the fast scan mode.

#### Parts for 3D Printing

- Sample chamber. You can download the CAD model as step file to send it to your 3D printing service.

#### Screw kit

You will need Hex Head Socket Cap metric screws:

| type | length (cm) | amount |
| --- | --- | --- |
| M3 | 10 | 4 |
| M4 | 30 | 4 |
| M4 | 25 | 2 |
| M4 | 20 | 5 |
| M4 | 10 | 2 |
| M5 | 30 | 2 |
| M6 | 20 | 3 |

E.g. this Srew kit will do it.

### Let’s Start to Assemble

#### The Light-Sheet Unit

Laser and Fiber Coupling

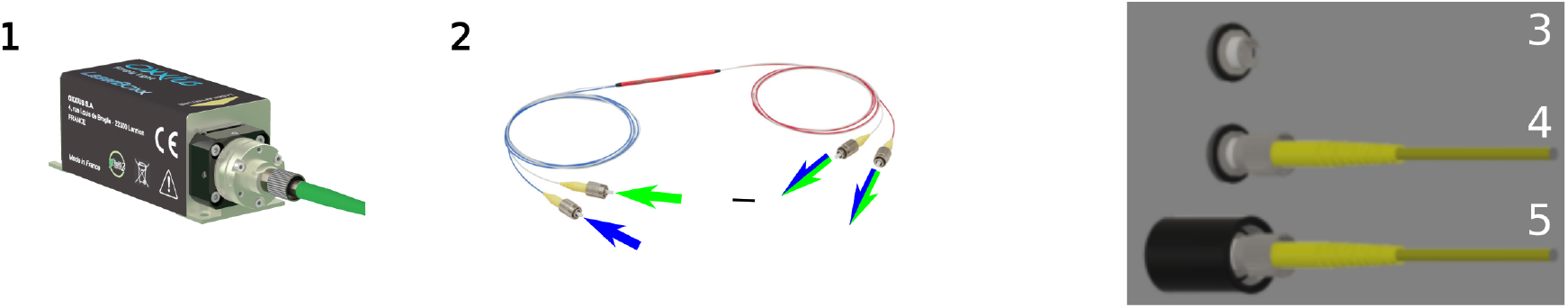

(1) Purchase continous lasers whose wavelength are adapted to your favorite fluorescence probes. We use a blue 488nm and a green 561nm laser. Choose the option with prealigned fiber coupler.
(2) To combine the two lasers into a single-mode fiber, attach the two inputs of the fiber optic coupler to the FC/PC connector of each laser. The blue and green lasers will be combined and available at both output fiber ports.
(3-5) Assemble the Fiber support tube: Take the FC/PC connector which will later hold the optical fiber that delivers the laser. Screw the connector into the lens tube (black) but not too far so that you can easily attach and detach the optical fiber. Fix the connector with the two retaining rings.

#### Adapt the Scientifica Scope to receive the light-sheet unit

You will need to unscrew the bracket that holds the objective support plate from the original Scientifica setup.

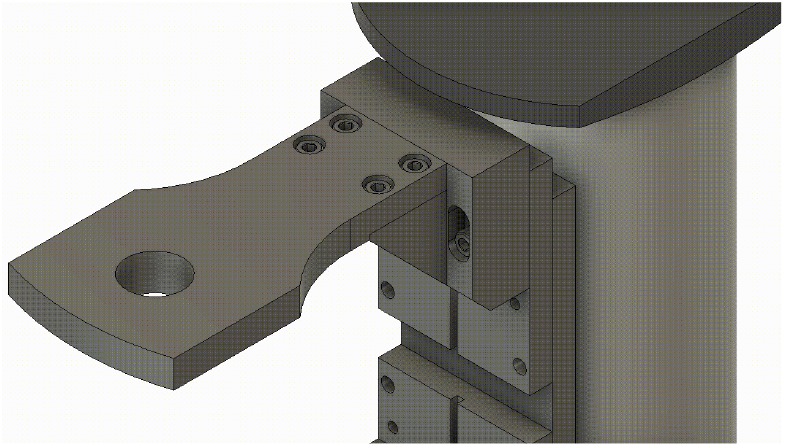

This bracket will then be replaced with a custom bracket that has holes to allow the light sheet unit to be attached to the Scientifica system in the next steps.

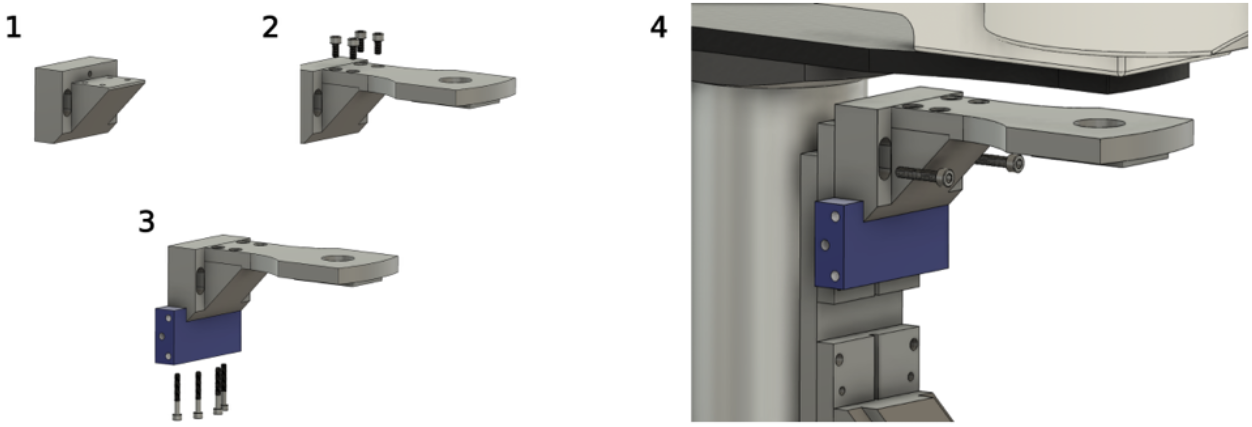

(1-2) Attach the top plate (previously unscrewed from the Scientifica Scope) to the Custom Bracket (custom piece) using the original screws.
(3) Attach the Adapter Plate 1 (custom part) with four M4×30 screws.
(4) Attach the assembly to the topmost Z-stage of the microscope with two original screws.

##### Assemble the light-sheet unit

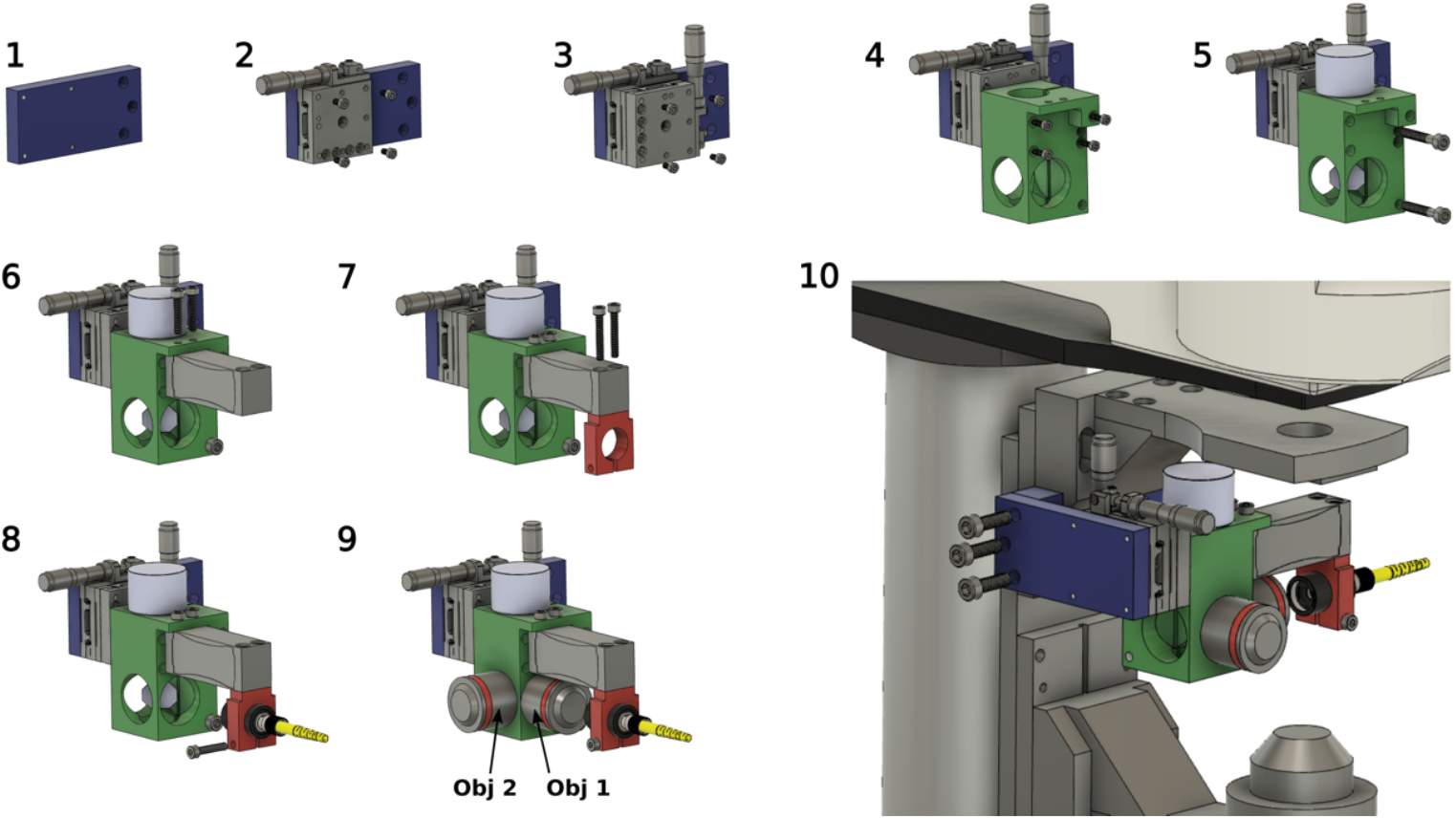

(1) Start from the Adapter Plate 2 (blue, custom piece).
(2) Attach the Horizontal Manual Stage to the Adapter Plate 2 with the four M3 screws included with the stage.
(3) Attach the Vertical Manual Stage to the Horizontal Manual Stage with the four M3 screws included with the stage.
(4) Attach the Light-Sheet Unit Central Cube (green, custom piece) to the Vertical Manual stage with four M3×10 screws.
(5) Insert the Galvanometer Mirror (light blue), align it at 45° relative to the sides of the Light-sheet Cube, and fasten it in place with two M5×30 screws.
(6) Attach the Piezocrystal (PZ 400 SG OEM). If you want to start with a simple low-cost version that uses only the slow scanning mode then install instead the Piezo Dummy (custom piece) to the Light-sheet Cube with two M4×20 screws.
(7) Attach the Fiber Holder (red, custom piece) to the Piezo with two M4×25 screws.
(8) Insert the Fiber support tube (see points 3-5) into the Fiber Holder, fasten it with one M4×20 screw and attach one of the output fibers of the fiber optic coupler to the connector (2).
(9) Screw in the Zeiss Plan-Neofluar Objectives (5x/0.16 M27). If your ultimate goal is to build the two-photon version then you should use instead the Olympus LMPLN5xIR/0.1 objectives, which is optimized for near-infrared transmission.
(10) Attach the Light-sheet Unit to the Scientifica Scope by fixing the adapter Plate 2 onto the Adapter Plate 1 using three M6×20 screws.

##### Laser Alignement

Before turning on the laser, prepare the room for laser safety (remove any jewelry, make sure no one is in the path of the laser, and that the laser does not hit any reflective surfaces). Unscrew the focusing objective (Obj 2) of the light-sheet unit. Switch on the galvanometric mirror and then switch on the laser at low power. By moving the fiber support tube (black) relative to the fiber holder (red), adjust the distance of the fiber outlet to the collimation objective (Obj 1) until the beam exiting the cube is well collimated.

Next, carefully center the beam in the opening where Obj 2 will be located by adjusting the orientation of the galvanometer mirror. Once this is done, secure the mirror position with the fixing screw. Secure Obj 2 in place again by screwing it in.

#### The Detection Path

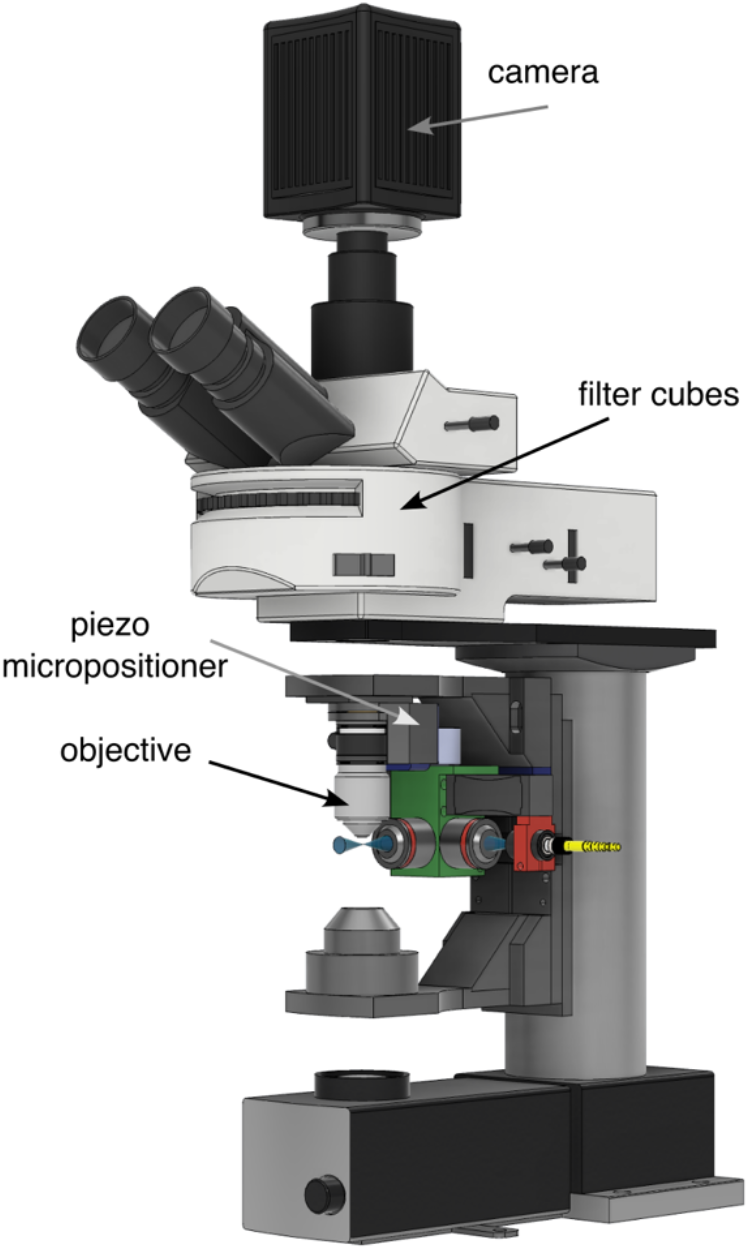

- To configure the detection path, use the standard ports and configuration of the Scientifica scope.
- Use a detection objective that is best adapted to your application. If you want to install the fast z-scan mode then attach the objective via an objective scanning piezo system.
- Position the filters into the filter wheel adapted to your application. For GFP imaging:
  ∘ a notch filter to block the 488nm laser line
  ∘ a GFP Emission Filter.
- Attach the camera to the camera port. In our systel, we use the Hamamatsu ORCA-Flash4.0 V3 camera

#### Driving the Light-Sheet Unit

The light-sheet unit can be operated in two modes:

##### Mode 1: Stand-Alone Light-Sheet Source Without the Need for Additional Computer Interfacing

To drive the galvanometer, a simple and affordable function generator is sufficient. Set the mirror to oscillate at approximately 400Hz with a triangular or sawtooth waveform.

If your image acquisition software allows the acquisition of z-stacks by controlling either the motorized stage of the z-focus or the piezo objective scanner, then you can operate the light-sheet unit in two modes.

##### Slow Scan Mode

Even if you have not installed the piezo-control of the objective and optical fiber, you can still perform volumetric recordings. Simply control the movement of the objective using the motorized stage of the Scientifica scope, similar to a standard epifluorescence recording. Since the light-sheet unit is attached to the same stage as the objective, it will move along with the objective, ensuring that the light-sheet always illuminates the focal plane.

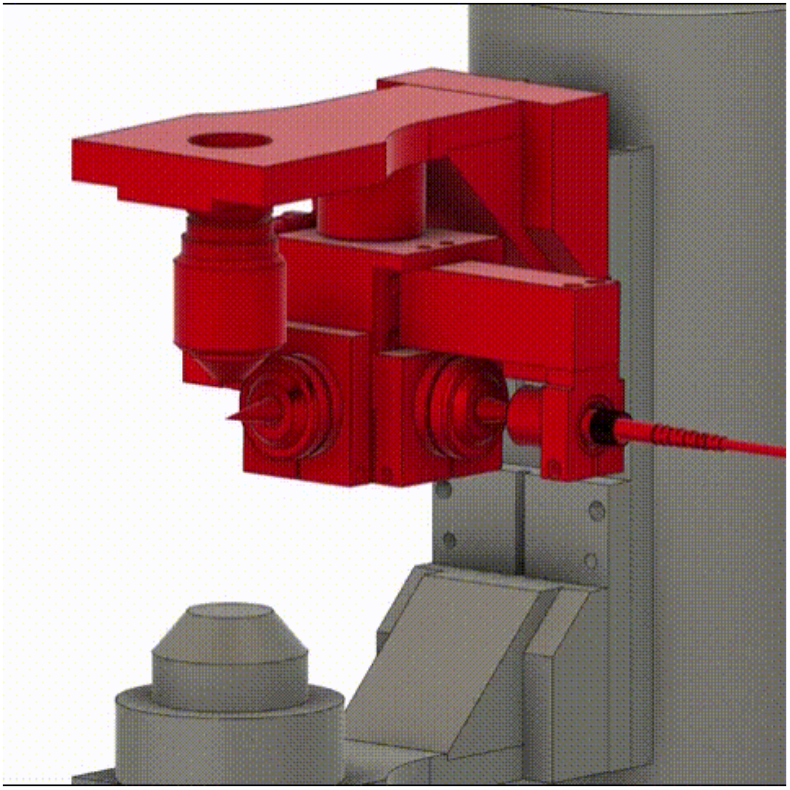

##### Mode 2: A Computer-Controlled Light-Sheet Scanner

###### Fast Scan Mode

If you have installed a piezoactuator for both the imaging objective and the fiber, you can achieve high precision and fast volumetric recordings. Synchronizing the movement of the fiber with that of the objective is crucial to ensure that the light sheet remains in the focal plane of the objective throughout the volumetric acquisition.

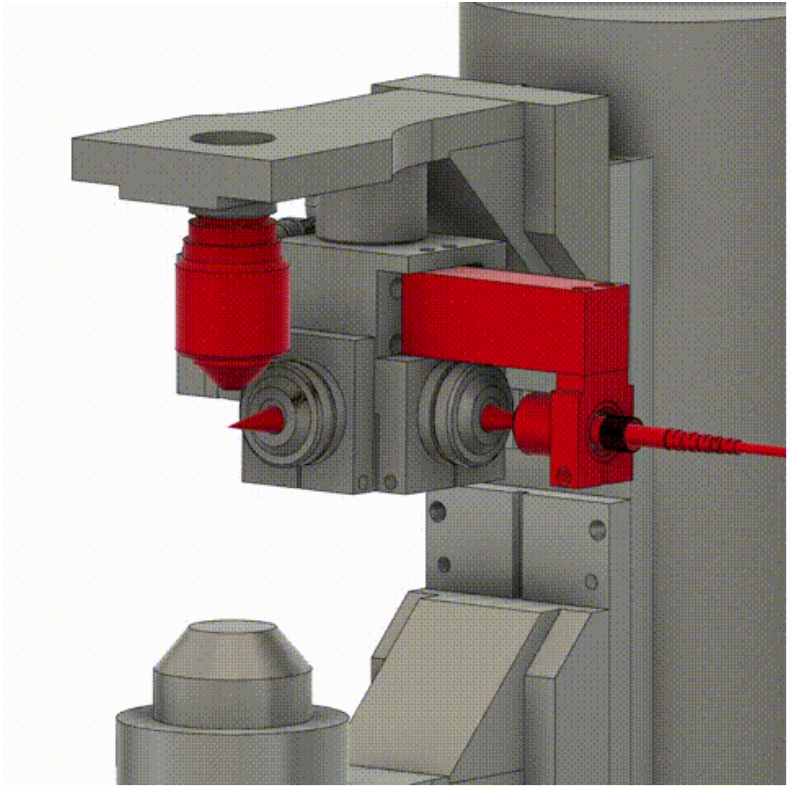

- To control the light-sheet volumetric scanning you can use our custom developed control software written in Matlab. Via a national instrument card PCIe-6363, this software simultaneously drives the objective piezo scanner, the galvomirror and the piezo that moves the fiber in order to scan the light-sheet through the sample. The software is open source.

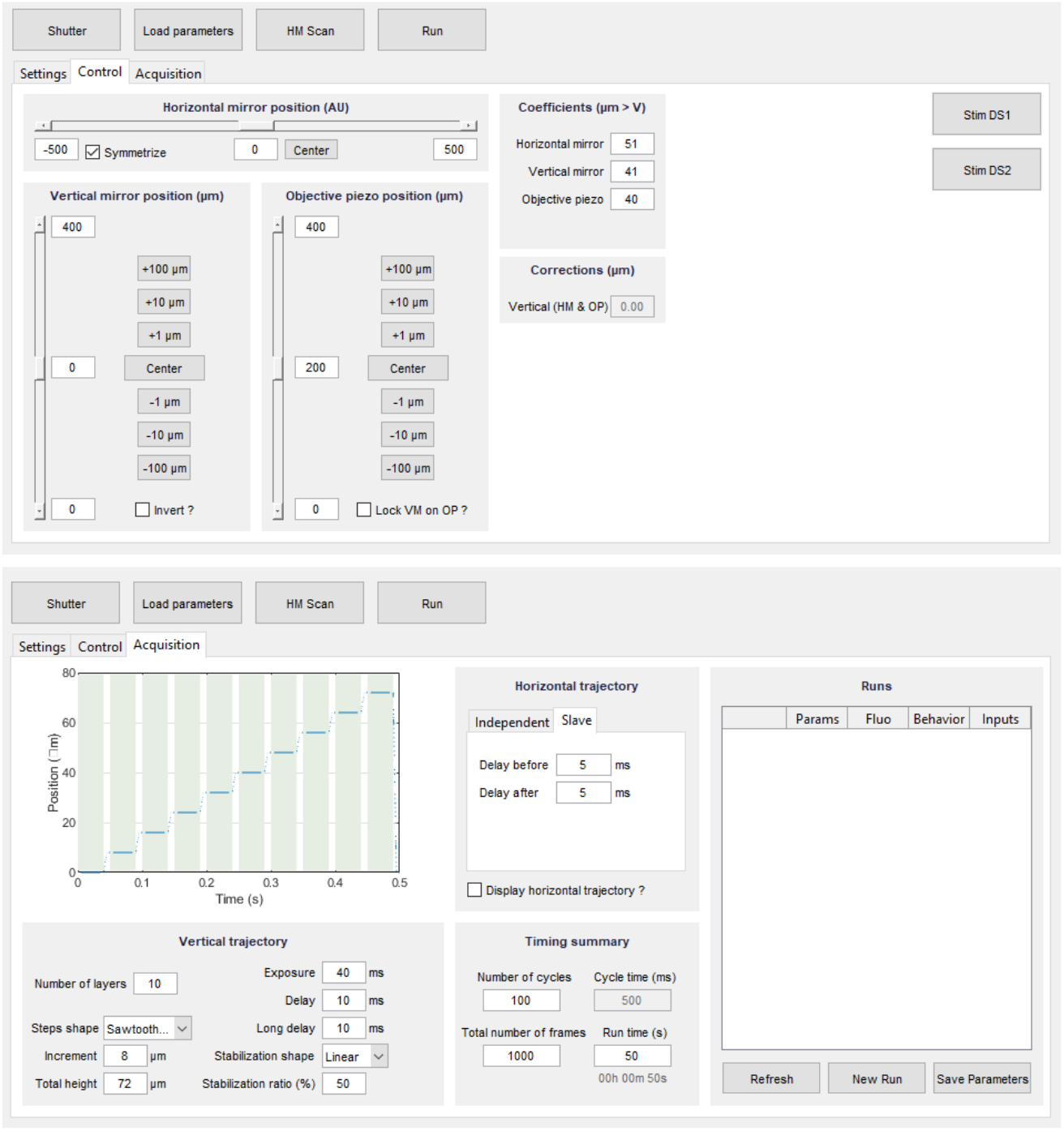

- Alternatively, you can use:
  ∘ ScanImage, the control software provided by Scientifica
  ∘ Micromanager, which supports the control of all Scientifica stages as well as of most common objective scanners from e.g. Physical Instrument or Piezo Jena.
  ∘ or the open-source python software presented here: https://www.frontiersin.org/articles/10.3389/fcell.2022.875044/full

#### The Sample Chamber Holder and Sample Chamber

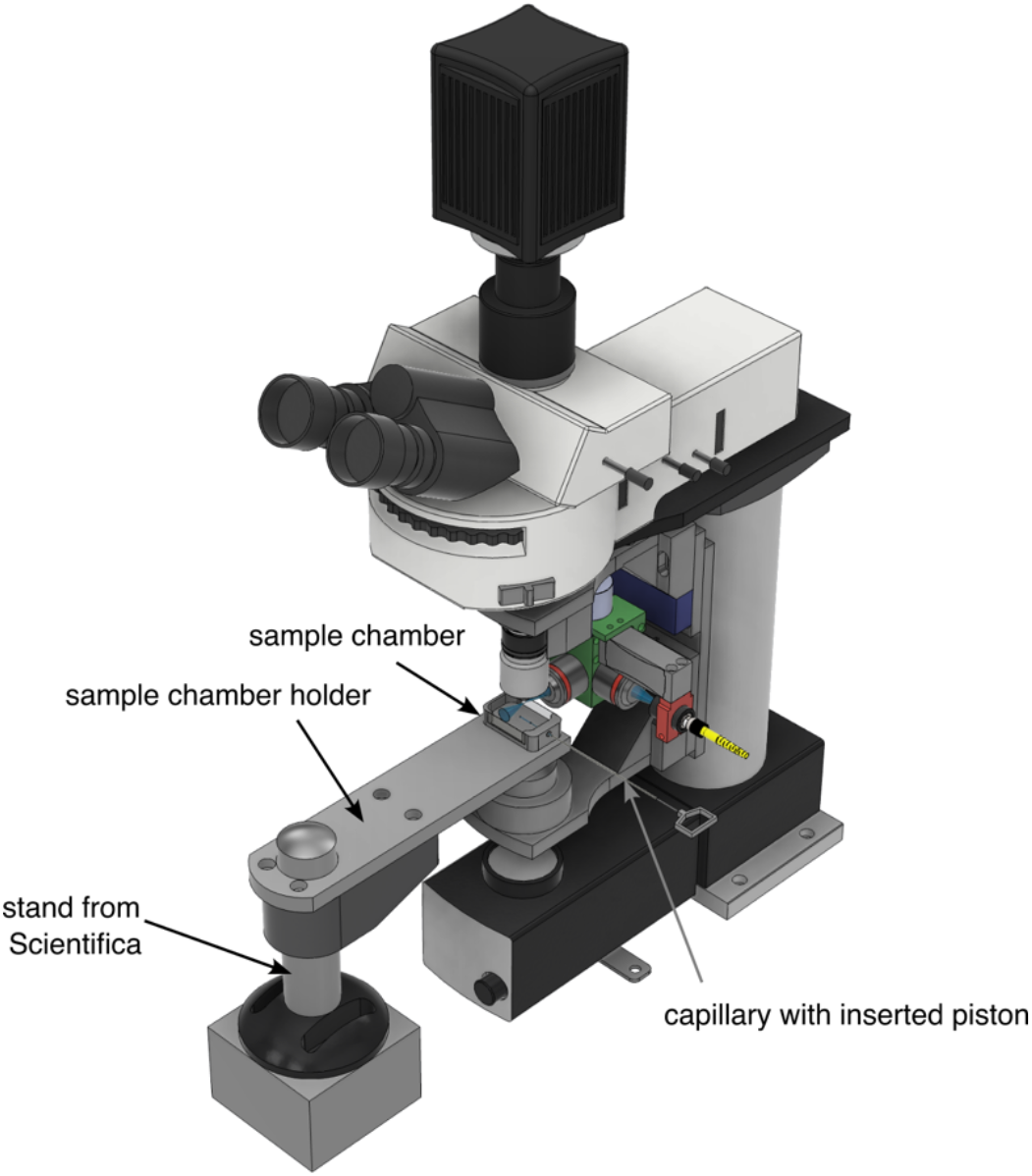

- Install the sample chamber holder (custom part).
- 3D print the sample chamber.

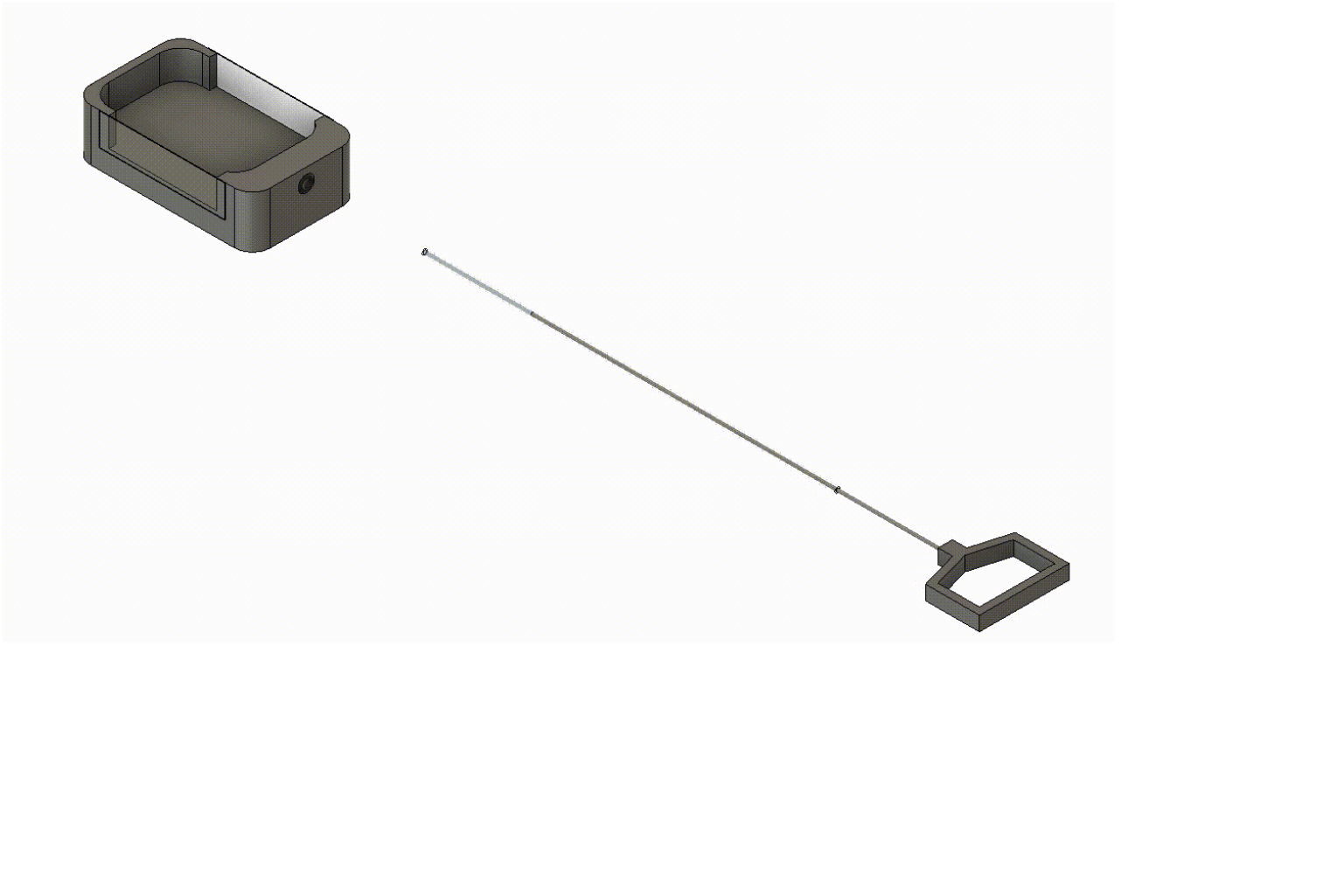

- Take the 3D printed sample chamber and use cyanoacrylate or UV-curing adhesive to glue a glass window on each side as well as an O-ring into the hole on the short side, which will allow you later to introduce and position the sample via a capillary (inner diameter 0.85mm, outer diameter 1.47mm, length 115mm) for the imaging sessions.
- Place the sample chamber into the predefined opening of the sample chamber holder.

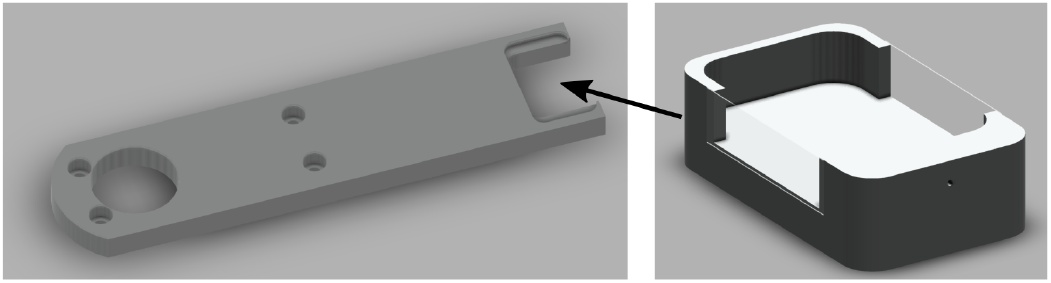

#### Ensuring Laser Safety

Protect the experimenter from the laser by installing a beam blocker in front of the sample chamber. Fix the T-bracket (custom piece) with two M4×20 screws to the sample chamber holder and then fix the beam blocker with two M4×10 screws to the T-bracket.

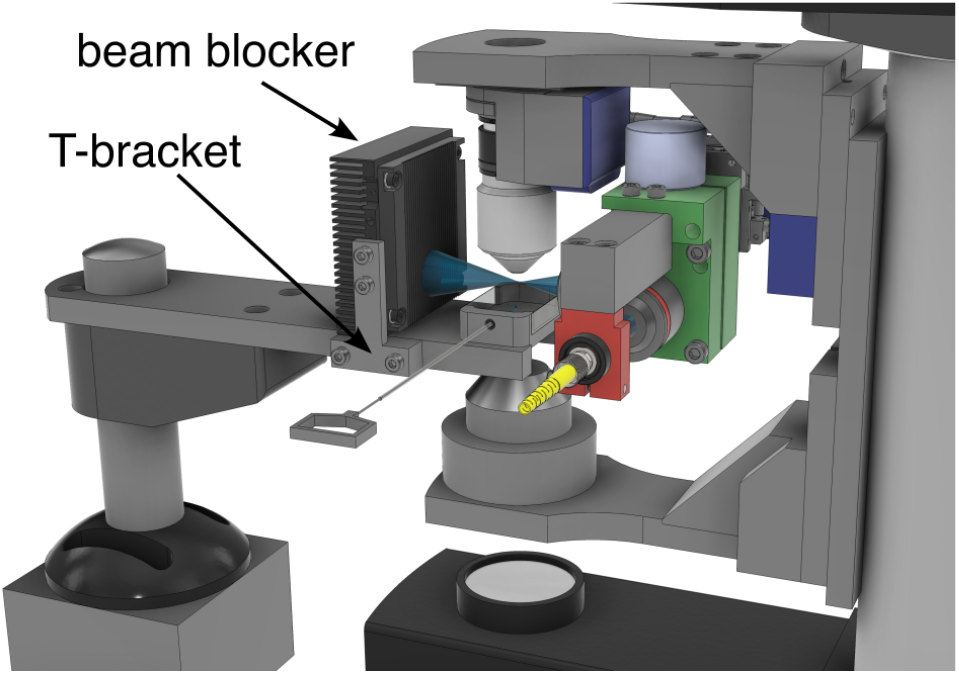

#### Now Let’s Align the System

- Fill the sample chamber with water and add a few drops of a 1.5mM fluorescein solution into the water-filled sample chamber to visualize the laser. Switch on the laser at low power until you can see the fluorescence laser profile. Be careful not to look directly into the laser and to keep working at low power (<1mW after the second illumination objective) for alignment.
- The use of fluorescein requires cleaning with ethanol or isopropanol after alignment to remove residual fluorophores that may cause imaging noise.
- It is also possible to use an agarose cylinder and to observe the scattered light from the laser, to do this remove the filter between the detection objective and the camera and follow the same procedure.
- Align the laser waist under the detection objective by moving the entire light-sheet forming unit with the x-translation stage
- Switch on the camera and align the laser into the focal plane of the detection objective by moving the light-sheet forming unit with the z-translation stage until you see a sharp image of the laser beam with the camera. If you do not see the laser beam, move the galvanometer mirror to bring it within the field of view.

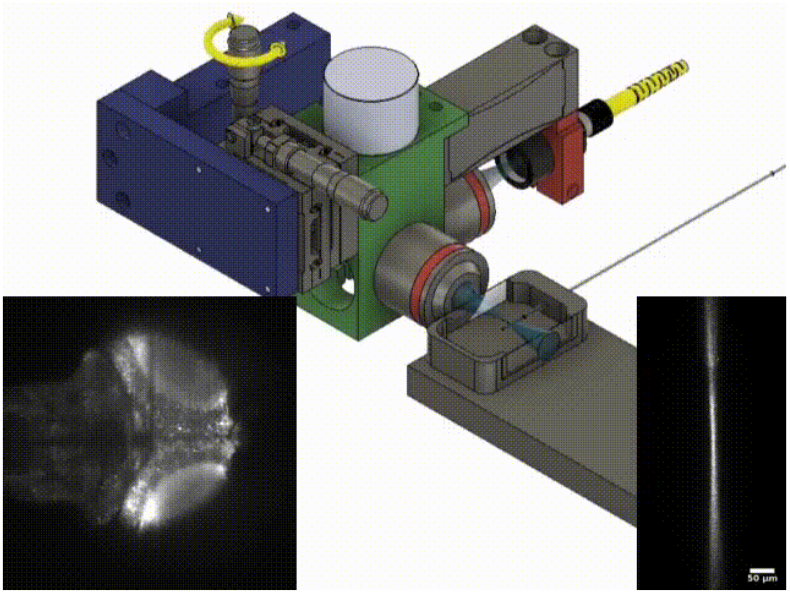

- Precisely align the laser waist in the center of the field of view using the x-translation stage

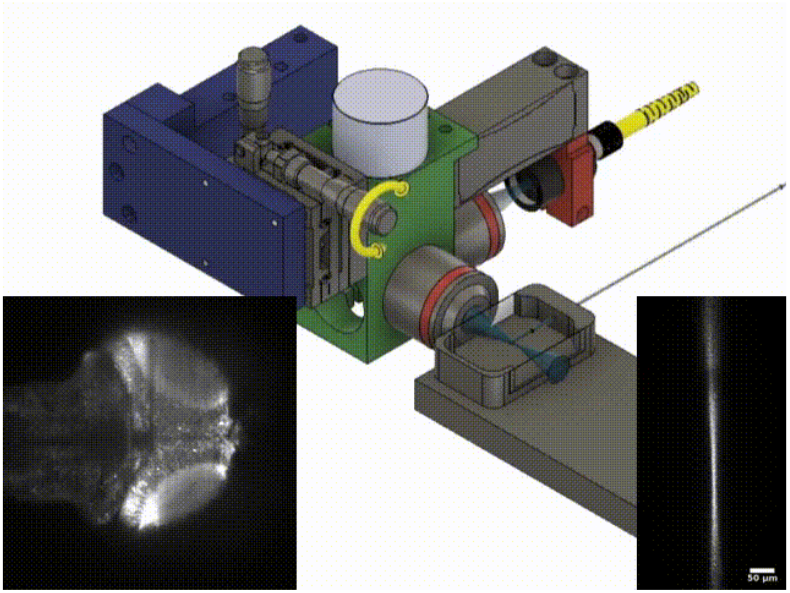

- Remove the fluorescein solution from the sample chamber and clean it.
- Now generate the light sheet by driving the galvanometer with a saw tooth pattern either with a function generator or via the control software (see above). Adjust the amplitude of the movement such that the light sheet covers the field of view.
- Now you can place your preferred sample into the light sheet and image it. Fine-tune the z-position and x-position of the light-sheet until you get the sharpest image possible.
- you can now produce 3D scans of your sample by manually moving the objective focus of the microscope or by recording 3D time-lapse movies using your microscope control software.
- Troubleshooting: if your image is not uniformly sharp throughout the field of view, the light sheet might be tilted with respect to the focal plane of the detection objective. In this case, you can correct this by tilting slightly the entire unit. For this, untighten the screws that connect the unit via the adapter plate to the microscope translation stage. Use a sheet of paper as a spacer between the adapter plate and the stage to tilt the unit in the correct direction once the screws are tighten. Repeat this procedure until your image is in focus across the entire field of view.

### Pictures of the System in Combination with an Electrophysiology System

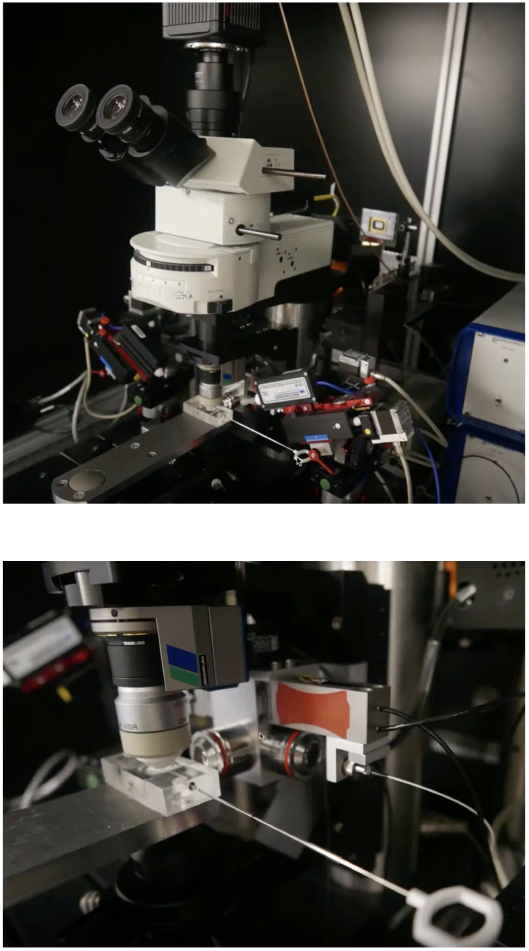

#### Example Recordings

Using the green laser. Shown is a high-resolution recording of a zebrafish brain (6dpf) with a pan-neuronally expressed red calcium indicator (elav3-jRGECO):

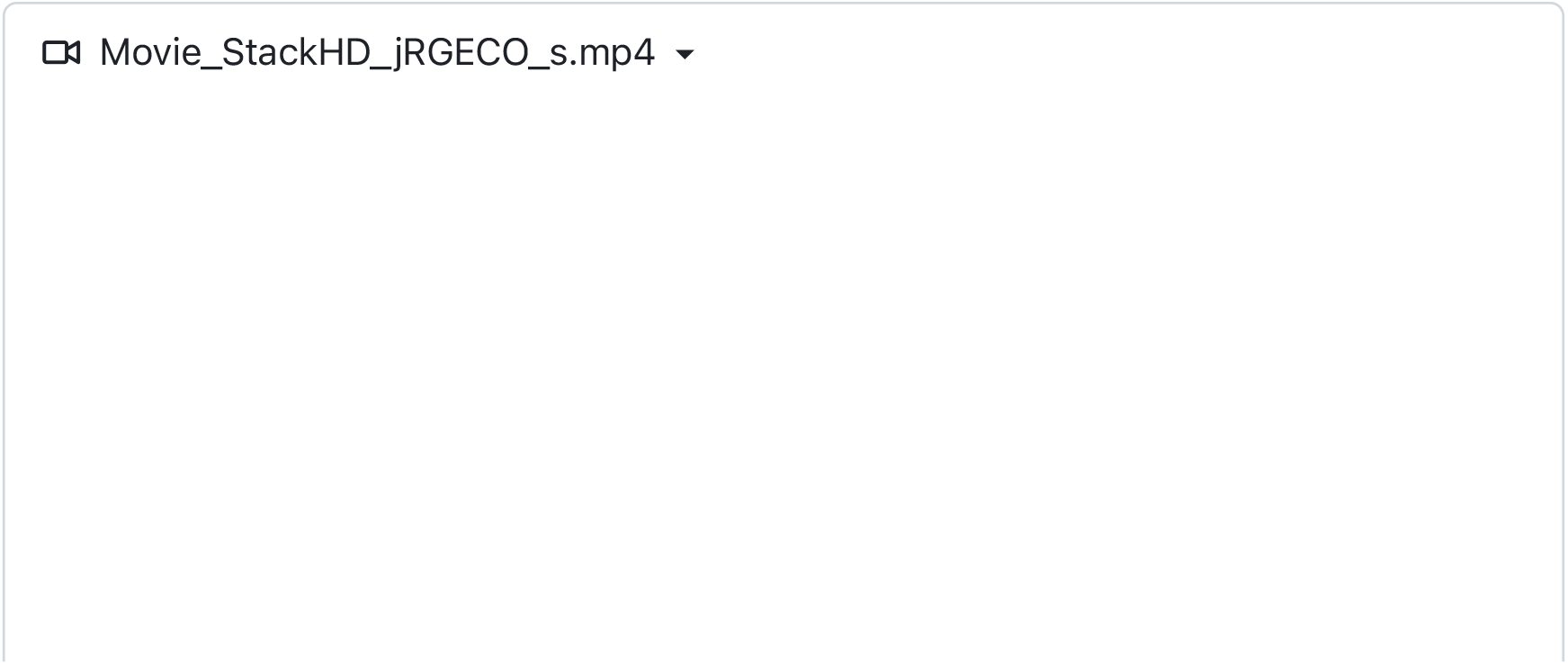
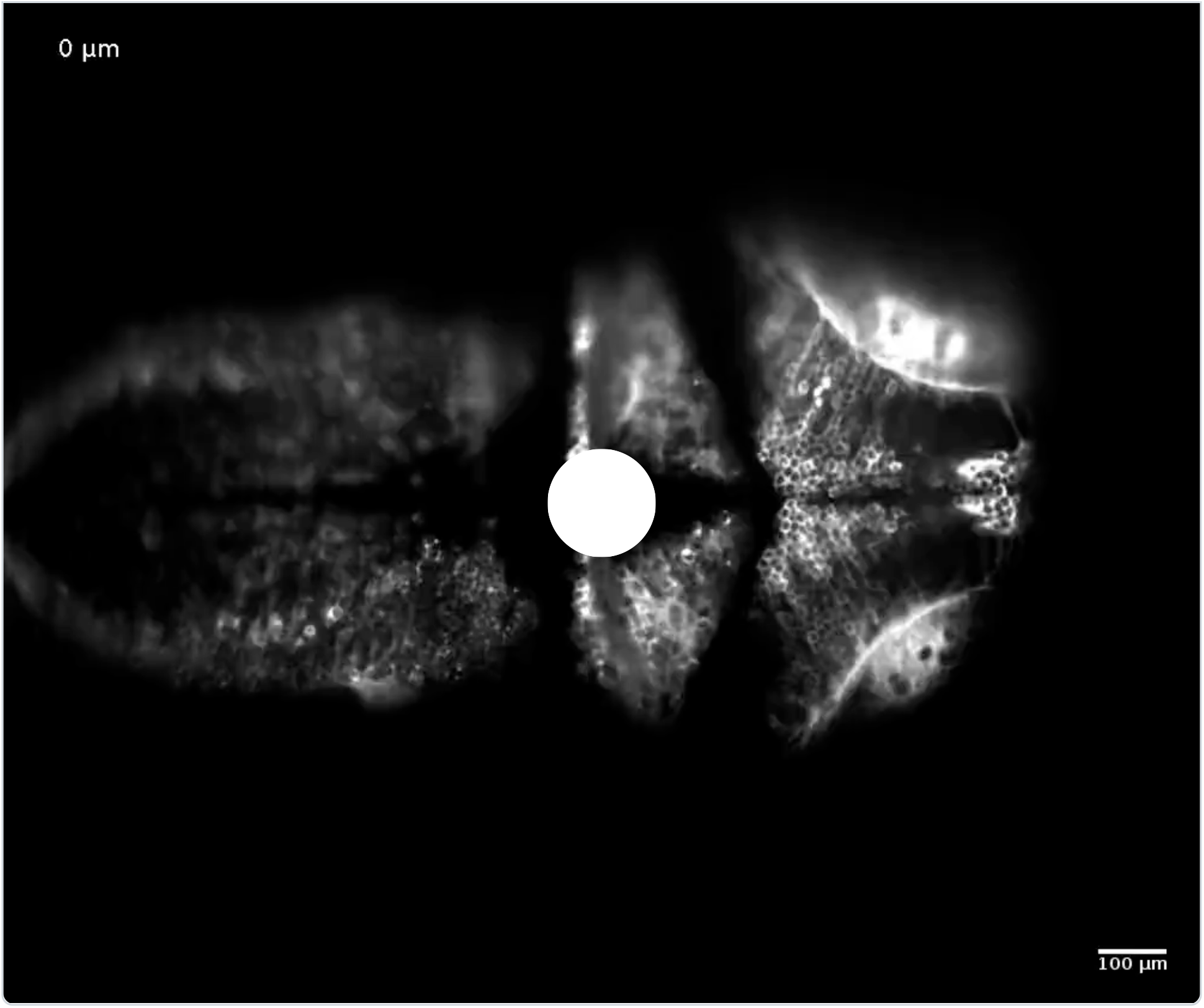

#### A Conference Talk Presenting the System

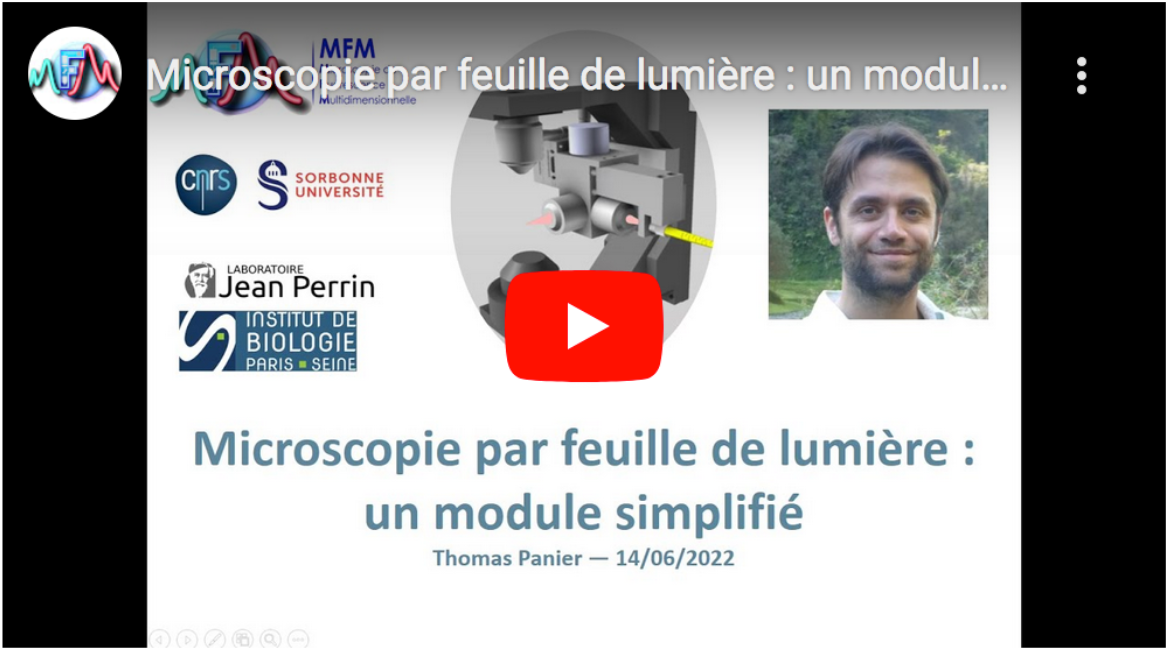

#### Upgrade the One-Photon Unit into a Two-Photon System by using a Hollowcore Crystal Fiber

Click here for detailed instructions

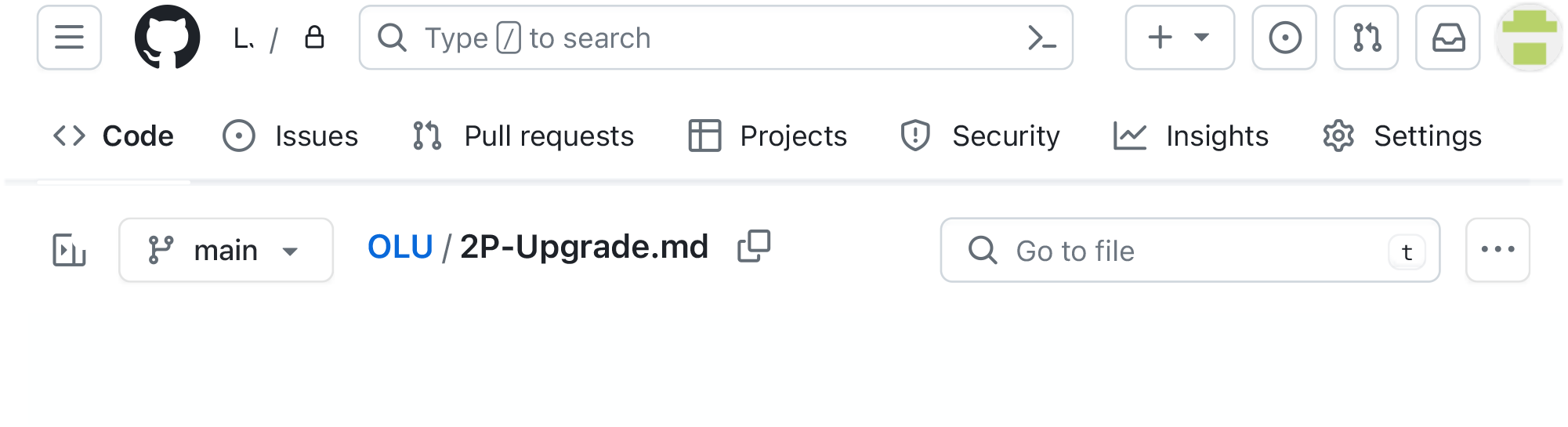

### Two-photon upgrade

This comprehensive guide provides detailed instructions for upgrading the system to the two-photon configuration. Assuming that you have already built the one-photon version of the system, the upgrade primarily involves two main steps. First, you need to fiber couple a two-photon femtosecond pulsed laser into a negative hollow-core optical fiber. Additionally, the fiber’s numerical aperture at the output side should be increased for compatibility with the light-sheet unit. Secondly, if not already implemented, the objectives used in the light-sheet unit must be replaced with objectives optimized for near-infrared transmission. Finally, a filter must be installed in the detection path to block the infrared laser. These modifications will enable efficient two-photon imaging with the system.

### Part List

List of parts

#### Custom Parts to Send for Milling

These parts are milled from an aluminum block. If you do not have a mechanical workshop in-house then you can send the *.step files that we provide below to an online milling service. For parts with threaded holes, join the mechanical drawings to the .step file.

- Custom part for compatibility with a MaiTai laser: View the 3D model or download the CAD model as a step file and the Mechanical drawing.

### Building Instructions

#### Multiphoton Light-Sheet Unit

The multiphoton light-sheet unit is basically identical in its design to the one-photon multicolor design. To build the two-photon system follow the building instruction of the 1P-Multicolor system but take into account the following modification:

For both the collimation and illumination objectives (step (9) of the building instruction of the Light-sheet Unit), use the Olympus LMPLN5xIR/0.1 objective optimized for near-infrared transmission. Although it is not optimized for visible wavelength, one can use this objective for one-photon imaging as well. To screw it into the light-sheet cube use a thread adaptor.

#### A Comprehensive Guide for Achieving Efficient Fiber Coupling of the Femtosecond Two-Photon Laser

##### Introducing Our Developed Negative Curvature Hollow Core Crystal Fiber

Together with GLO Photonics, we selected an HC-NCF fiber whose optical characteristics are ideally suited for laser delivery in the context of combined one- and two-photon light-sheet microscopy.

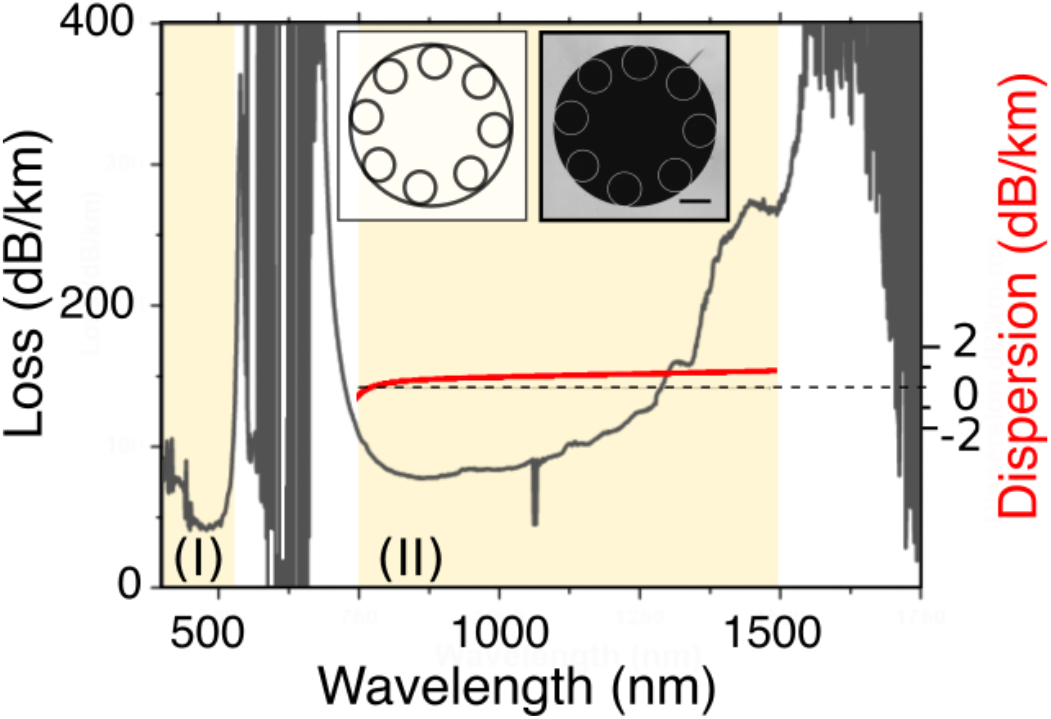

The outer fiber diameter is 200 ± 2 µm and the fiber coating diameter is 600 ± 30 µm. The fiber has an air-filled 30 µm-in-diameter core surrounded by a ring of eight non-contacting ∼10 µm-in-diameter cladding tubes. In this so-called hypocycloid fiber structure, the negative curvature of the core contour is created by the tubes surrounding the core. Tube diameters, inter-tube distance, tube wall thickness, and core size together control the fiber transmission spectrum. They yield two broad transmission bands that are compatible with 1, 2, and 3P imaging, respectively (see figure above): One in the near-infrared from 700 - 1500 nm (I), and another in the visible spectrum from 400 - 535 nm (II). Within these spectral bands, the loss is less than 300 dB/km (corresponding to a transmission efficiency of more than 93% for a one-meter-long fiber), and the pulse dispersion is less than 1 dB/(km·nm) of spectral pulse width.

The fiber is commercially available as a patch chord cable with standard FC/PC connectors. This proves extremely convenient as it allows one to disconnect and reconnect the fiber to the optical setup without the need for subsequent realignment on either end. The small core diameter compared to the infrared wavelength further ensures near single-mode guidance.

The Gaussian beam properties of the laser are also well preserved at the fiber output with M^2 values of 1.23 and 1.18, measured at 1030 nm and 515 nm laser wavelength respectively. The near-field and far-field profiles are symmetric with minimal ellipticity of 97.25% and 85.57%. The mode field diameters at 1/e^2 are 23 ± 1 µm and 26 ± 1 µm, respectively, which correspond to a fiber’s numerical aperture (NA) of approximately 0.02. Due to negligible interactions between the light and the fiber material, the fiber exhibits a very high damage threshold: we could not observe any fiber damage up to the maximal tested peak powers of 40 µJ at 1030 nm and 10 µJ at 515 nm for 400 fs pulses, corresponding to an average laser power of 20W and 4W, respectively. These characteristics demonstrate that high-quality Gaussian beams can be delivered with this fiber across a broad spectral range with high transmission and low dispersion, which is a prerequisite for its application in high-resolution microscopy.

##### The Optical Path for Laser-to-Fiber Coupling

The following schematic and 3D model that you can interactively explore show a possible optical path for the laser coupling. You can adapt the arrangement to the space availability on your setup (mirror, **M**, lens, L, dichroic, **D**, wave plate, **W**, polarizer, **P**, beam trap, **BT**):

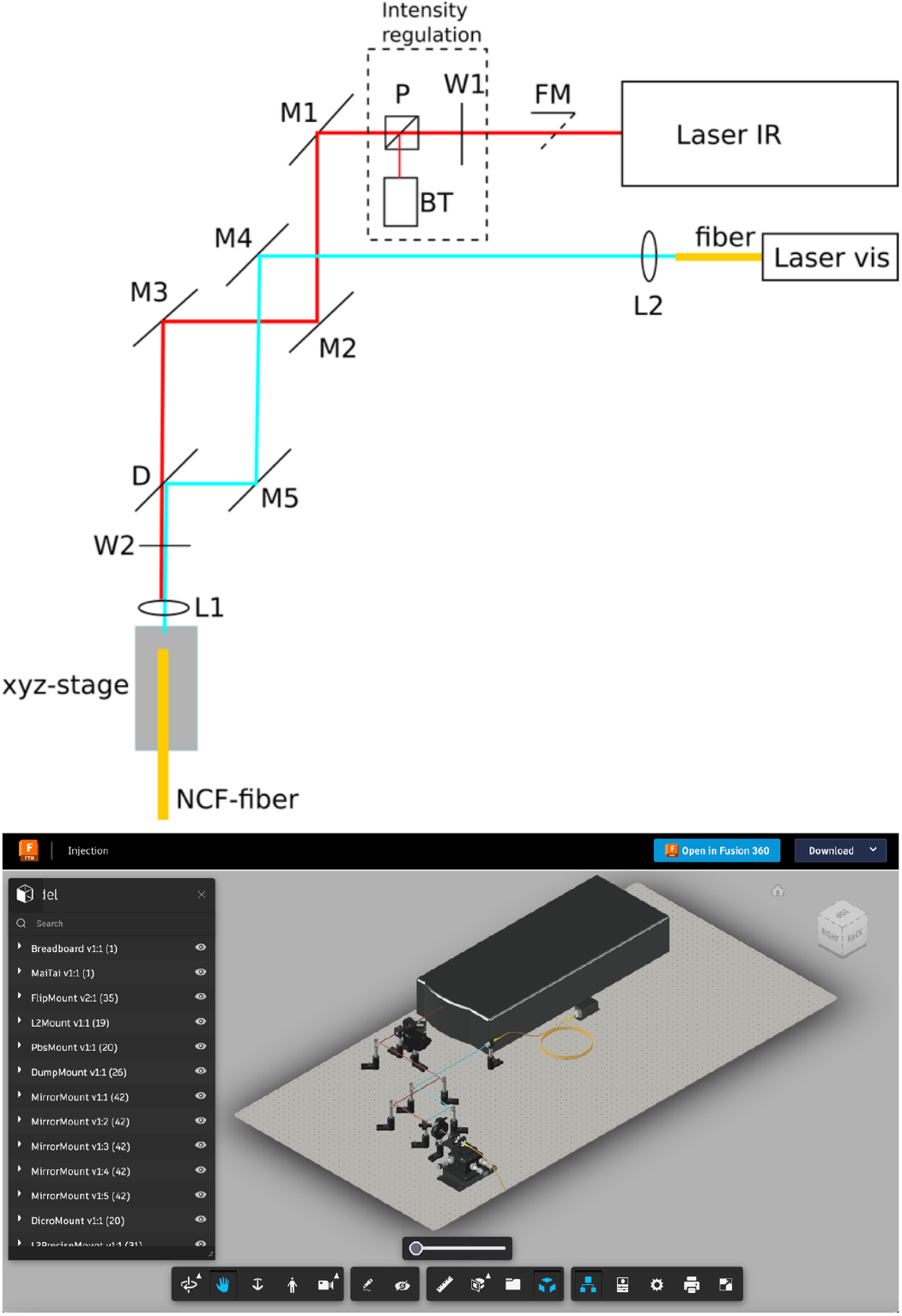

If you do not have space on your table you can mount a platform on top of your IR-pulsed laser source and bring the laser beam with a periscope up to the platform.

In the following, we propose a step-by-step explanation of how to build this optical path and how to align the optical components to achieve optimal fiber coupling. But before we begin, we will give some preparatory notes and general background:

##### Tips and Techniques for Laser Coupling into the Broadband Hollowcore Crystal Fiber

For the alignment we will follow a protocol well explained in an excellent YouTube tutorial that you find here and that we advise you to watch carefully before starting. The video explains a first pre-alignment step based on the backpropagation of a laser in the reverse direction. To use this trick first connect a standard single-mode fiber to the coupling unit. Use a fiber tester to inject a visible laser through the fiber. Then follow the steps as in the tutorial to co-align the alignment laser and the IR laser. Then disconnect the fiber and connect the hollow core fiber. If the first step was done correctly you should have enough transmission through the fiber to be able to further optimize the fiber coupling as described in the tutorial. To watch the video just click on the snapshot:

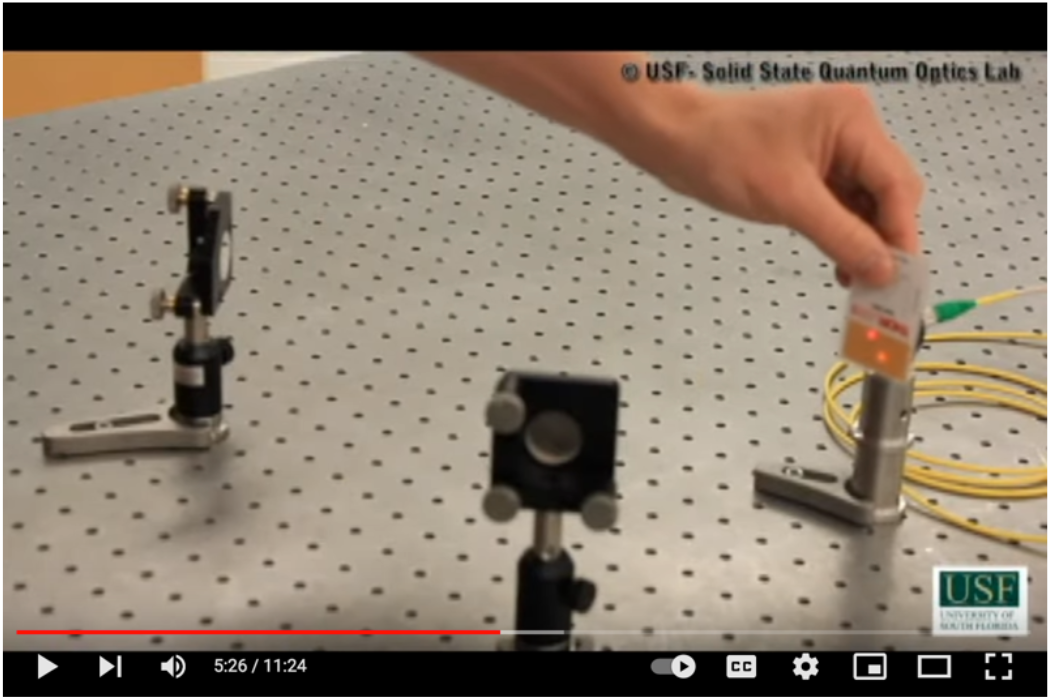

##### Mastering the Handling of Hollow Core Fibers: Tips and Techniques

- Here you find the fiber spec sheet of our hollow-core negative curvature fiber
- Attention !: The hollow core fiber front face is not protected and cannot be easily polished, which is a standard procedure used for conventional single-mode fibers. Work clean and remove dust before connecting the fiber to a porte. Also never try to clean the fiber outlet with ethanol or acetone because these solutions will enter the fiber core by capillary forces making the fiber unusable. You can use a fiber inspection scope to inspect the fiber core.
- The fiber has a very high damage threshold but only if the laser is well coupled. So it is advised to do the alignment procedure at low laser power. This improves also laser safety. You can control the laser intensity with a rotatable lambda wave plate installed in series with a polarizor.

##### Selecting the Optimal Coupling Lens: Criteria and Considerations

- For efficient optical coupling and suppression of higher laser modes, the diameter of the laser spot projected onto the fiber input side has to match the mode field diameter of the fiber. Or in other words, the opening angle of the focused laser beam has to match the numerical aperture of the fiber. Our fiber has a numerical aperture of 0.02 (see fiber spec sheet). The numerical aperture of the coupling system is given by 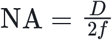, with ***D*** the diameter of the laser beam at the position of the coupling lens, and *f* the coupling lens’ focal length. The laser company normally gives the beam width and the beam divergent angle. In our case, the Ti:Sapphire laser (MaiTai, Coherent, USA) has an output beam waist < **1.2mm** and a divergent angle of < **0.001mrad**. We measured a beam diameter of 1.6mm at the position where we decided to install the coupling unit and we thus chose a coupling lens with focal length 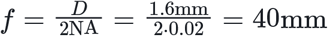. We got reasonable coupling with the achromatic coupling lens. If you want to further optimize the coupling you can use an objective as coupling lens. And you can install a telescope to adjust the beam diameter to match the fiber numerical aperture with the chosen coupling lens focal distance. Further tips for fiber coupling can be found in the manual from GLO-photonics.
- To measure the laser beam diameter, you may use either a beam profiler (Thorlabs BC207VIS/M) or the moving knife technique. For the latter, fix a razor plate on a linear translation stage. For this use a thin plate holder, a right angle clamp and a post system. Then move the knife perpendicular to the laser path step-by-step out of the laser beam while measuring the laser intensity as a function of the knife-edge position. The measured normalized power as a function of knife-edge position, *x*, can be fitted by 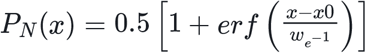. The fit parameter 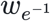 is the beam radius at *e*^−1^. You can also read 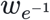 from the graph as half of the distance between the positions where the normalized power has a value between 0.08 and 0.92. Note that the beam diameter, ***D***, relevant to calculate the numerical aperture is measured at *e*^−2^ and is thus given by 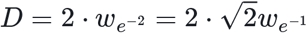

#### Let’s Get Started: Building the System

##### Coupling the IR Laser into the Broadband Fiber

- Position the motorized flip mount (FM) holding a protected silver mirror directly after the infrared laser output to hijack the laser for coupling into the optical fiber. Flipping the mirror allows you to select the setup into which you want to direct the laser. Alternatively, you can use a beam splitter to use the laser source on both setups simultaneously. The pertinence of the latter solution depends on the laser power that you need in both setups for your experiments. For a standard scanning two-photon system, you will generally need less than 100mW. A MaiTai laser has an average output power of 1.8W at 915nm wavelength. With a 90:10% beam splitter and assuming 50% loss of laser power until the sample you can perform light-sheet microscopy with an average laser power of 400mW and two-photon scanning on the standard system with a mean power of 90mW.
- Next install the optics to control the laser intensity. Position the lambda wave plate and the polarizor in series.
- Before switching on the laser, prepare for laser safety in the room and wear laser safety glasses to protect your eyes!
- Install the power meter to measure the laser power after the polarizer (**P**)
- Switch on the laser and rotate the wave plate until the transmitted laser power is minimal.
- Increase again slowly the laser power until you start seeing the laser spot on the IR detection card
- Close the laser shutter
- Position the two protected silver mirrors (**M1, M2 & M3**) mounted in kinematic mirror mounts. With mirror **M2** and **M3** you will later align the laser with the fiber axis. A good distance between these two mirrors is about 20cm. Adjust the height of the posts that hold the mirrors with the height of your laser beam. Tighten them on the optical table with Universal Post Holders.
- Install the long pass dichroic mirror (**D**) that is transparent for the infrared laser but reflects the visible spectrum. This will allow later the coupling of an additional visible laser into the same fiber.
- Mount the differential **xyz-translation stage** (MAX313D/M) at about 20cm from the mirror **M3**. With this stage, you will position the fiber outlet precisely into the focal point of the coupling lens. You might have to mount this stage on a platform to match the beam height (**Custom piece for compatibility with a MaiTai laser:** Mechanical drawing, .stl file **and** .step file). Alternatively, you can lower the beam path with a periscope.
- Install the half-wave plate (W2) mounted in a high-precision rotation mount to adjust the polarization of the laser. For the highest signal in two-photon microscopy, the laser polarisation needs to be aligned with the light-sheet plane. You can do this later by rotating this waveplate until you observe a maximum of the fluorescence signal.
- Fix the mounting bracket onto the non-moving front of the **xyz-stage** and mount onto it a SM1-Compatible Flexure Stage Mount to hold the coupling lens **L1**.
- Screw the coupling lens into the holder (AC254-040-B-ML)
- Mount the SM1-compatible flexure stage mount onto the moving platform of the xyz-stage and screw into this mount the FC/PC fiber adapter plate with its external SM1 threading. This FC/PC adapter will later receive the optical fiber.
- Attach the single mode fiber for prealignment to the FC/PC adaptor plate.
- Attach the other side of the fiber to the cable continuity tester for pre-alignment.
- Switch on the red laser of the continuity tester. You should see the laser exiting out of the other side of the fiber
- Adjust the distance of the coupling lens (**L1**) relative to the fiber outlet until the red laser light from the continuity tester is well collimated by the coupling lens.
- Now follow the alignment steps explained in the YouTube tutorial. Eventually, watch the video again.
- Now that you know how to do it ensure laser safety in the room and wear laser safety glasses to protect your eyes!
- Prepare a VIS/IR Detector Card to visualize the invisible infrared laser Open the laser shutter.
- Now move the two mirrors **M2** and **M3** to co-align the red laser and the IR laser as was explained in the video.
- Switch off the laser shutter
- Detach the single-mode fiber from the coupling unit
- Attach the negative curvature broadband hollow core fiber to the coupling unit. Position the power meter before the coupling lens and adjust the power to about ??? mW.
- Install an FC/PC adaptor plate and fix to it the other end of the fiber. Install the power meter just after the fiber to measure the transmitted laser power.
- Open again the laser shutter. You should measure laser transmission through the fiber.
- Now adjust the mirrors **M2** and **M3** as was described in the video until you get > 90% laser transmission.
- Close the laser shutter

##### Coupling the Visible Laser into the Broadband Fiber

- Fix the fiber coupled laser onto the table.
- Position mirror **M3** and **M4** with a distance of about 20cm between each other such that mirror **M4** is placed about 20cm from the coupling lens. As mirrors choose again protected silver mirrors mounted in kinematic mirror mounts to align the laser with the fiber axis. Select the length of the posts compatible with the height of your laser beam.
- Attach the fiber splitter to the laser that you purchased with the prealigned fiber coupler.
- Install a prealigned fiber collimators **L2** that you mount into a kinematic mirror mount using an adaptor and positioned onto the adequate posts and its universal post holder.
  ∘ We selected the collimation lens focal distance with the formula: 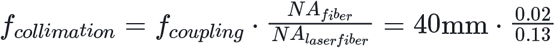 which boils down to the fact to choose the collimation lens focal distance such that the collimated laser beam diameter matches the diameter of the infrared beam that we previously coupled into the fiber using the chosen coupling lens.
- Attach one of the outputs of the fiber coupler to this collimator lens.
- Attach the other side to the FC/PC connector plate on the **xyz-stage** of the coupling unit. If the fiber is to short you can extend it with a fiber-to-fiber connector and the single mode fiber that you have used in the previous pre-alignment step.
- prepare laser safety and switch on the laser at the lowest power possible where you can see with the VIS/IR Detector Card the laser.
- Now use the mirror **M3** and **M4** to co-align the two laser beams that travel in the reverse direction through the system as explained in the alignment YouTube tutorial.
- Switch off the laser and replace the single-mode fiber that is connected to the coupling unit with the negative curvature fiber.
- Switch the laser again on.
- You should detect with the power meter a transmission through the fiber. Adjust mirror **M3** and **M4** until you have more than 75% percent of transmission

At 915nm wavelength, the central two-photon absorption peak of GFP and of its calcium sensitive derivative GCaMP, we delivered through 1.5m fiber length 100fs laser pulses with 98% power transmission efficiency and minute pulse dispersion of 28nm (1dB/km·nm) that we fully precompensated with the Deepsee element of the MaiTai laser source. At 488nm, the one-photon excitation maximum of GFP and GCaMP, we achieved a transmission efficiency of ∼75%.

Go back to the main page

#### Increasing the Fiber Numerical Aperture for Compatibility with the Light-Sheet Unit

- The collimation and illumination objective form a one-to-one telescope. If we would simply connect the hollow core fiber to the light-sheet unit as we did in the 1P-Multicolor system then the light-sheet thickness would correspond to the mean field diameter of the hollow core fiber which is **23*µm*** (fiber spec sheet). To get the same resolution as in the one-photon implementation we have to install an extra lens after the fiber to demagnify the laser waist from the fiber output to about 5*µm* and at the same time to increase the divergence angle of the laser to match the numerical aperture of the collimation objective. The numerical aperture of the fiber is 0.02 and thus a factor 5 times smaller compared to the numerical aperture of the detection objective.
- With a lens with a focal distance of f = 2mm and a numerical aperture of 0.5 you can demagnify the laser output from the fiber by a magnification of m = 0.2. Mount this lens via an adaptor into the lens tube which you prepared in step (5-7) of the manual of the 1P-Multicolor unit.
- Place the lens at a distance 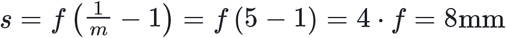 after the fiber output. The lens refocuses the laser at a new position 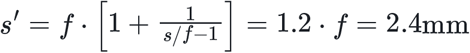 after the lens down to a waist of 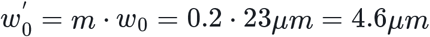 with an angle of divergence of the laser corresponding to a numerical aperture of ***m* · 0.02 = 0.1** thus matching the numerical aperture of the collimation objective.

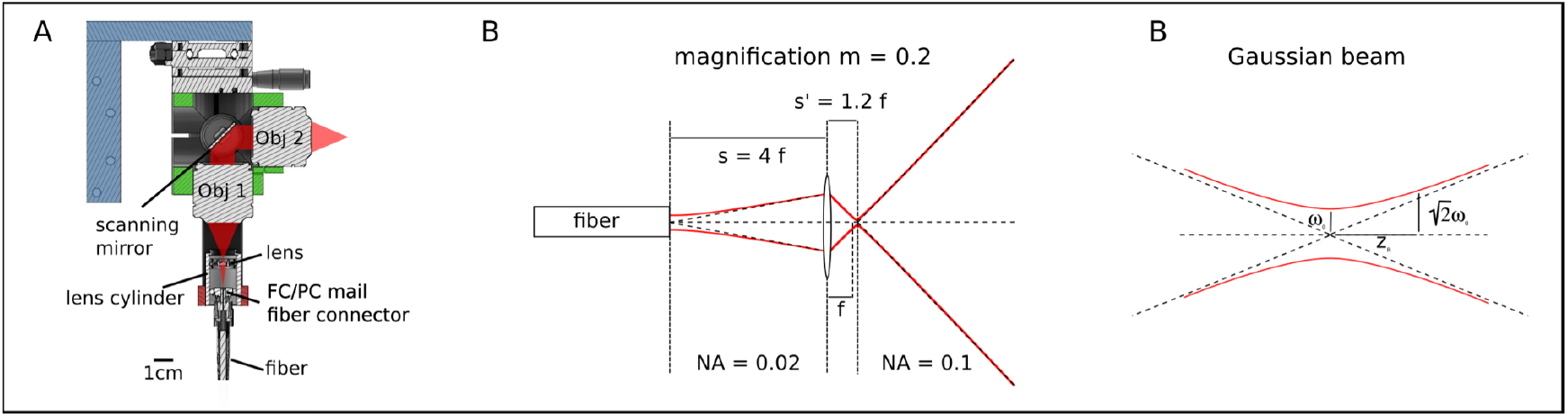

Focusing of spherical Gaussian beams by Sidney A. Self

#### Modifying the Fluorescence Detection Path to Block Infrared Illumination

- Install in addition the multiphoton short-pass emission filter to block also the pulsed infrared laser source.

#### Alignment of the setup

- Align the system in one-photon mode following the instructions provided for the 1P-Multiphoton system. This alignment procedure will also align the two-photon pathway.
- Align the polarization of the two-photon laser within the light-sheet plane. Visualize the two-photon laser using a fluorescein solution. Rotate wave plate W2, which is installed in front of the xyz-translation stage in the fiber coupling optical pathway. Continue rotating the wave plate until the fluorescence signal is maximized, indicating the optimal position.
- Note that the alignment and transmission efficiency of the hollow-core fiber in the infrared are not affected by fiber bendings. However, bending the fiber will indeed affect the polarization by changing its direction and ellipticity. Once you have set the rotation angle of wave plate W2, it is crucial not to move the fiber any further. If the fiber is inadvertently moved, the rotation angle of the wave plate will need to be readjusted accordingly to ensure optimal performance. For further information regarding the dependency of light-sheet fluorescence microscopy on polarization, please refer to Vito et al. Optic Express 2020.

### Example of High-Resolution Zebrafish Brain Recordings in One- and Two-Photon Mode (elav3:H2B-GCaMP6)

- Left: one-photon mode excited @ 488nm
- Right: two-photon mode excited @ 915nm

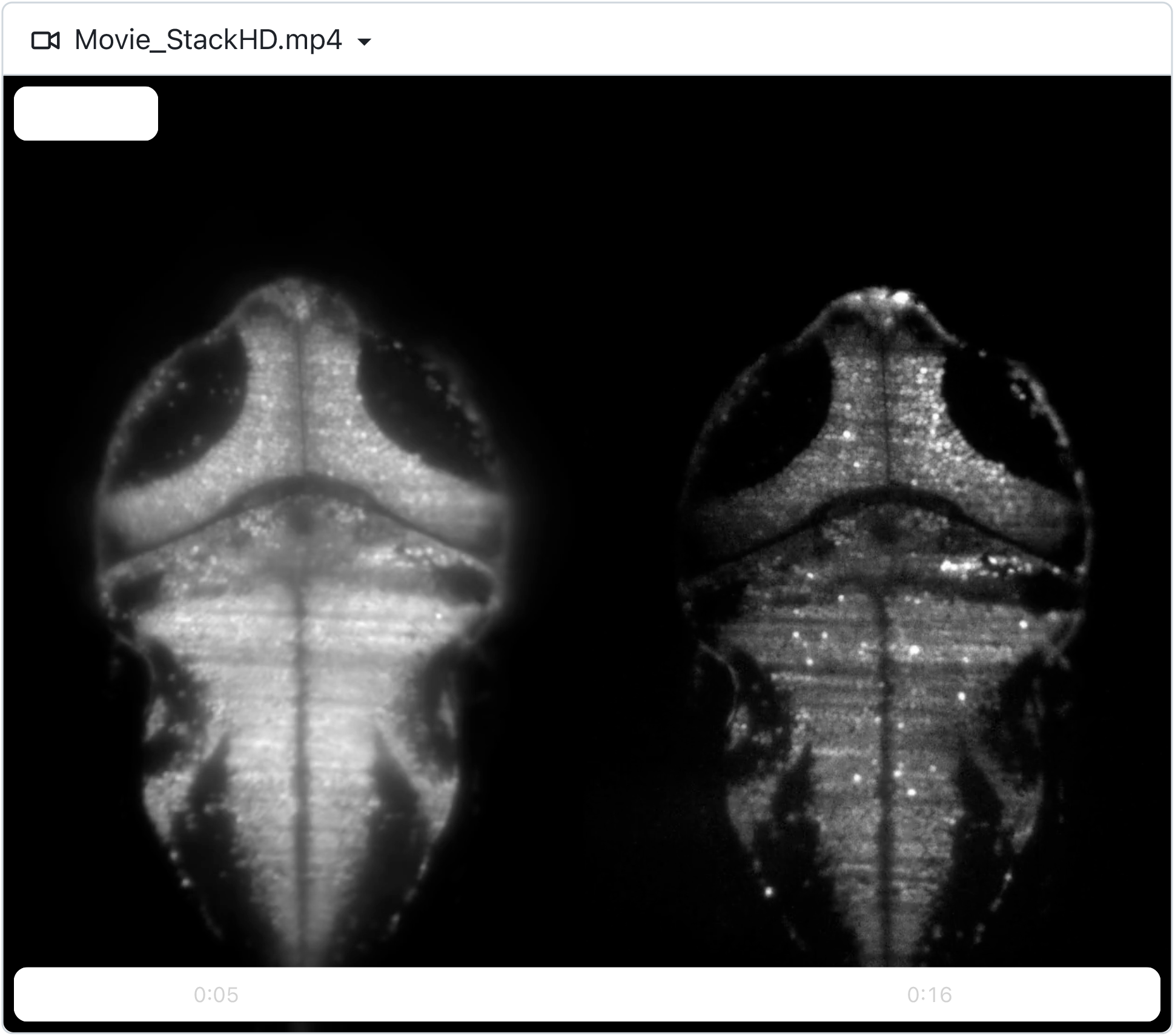

#### Mechanical drawings of custom parts

**Fig. S1.**
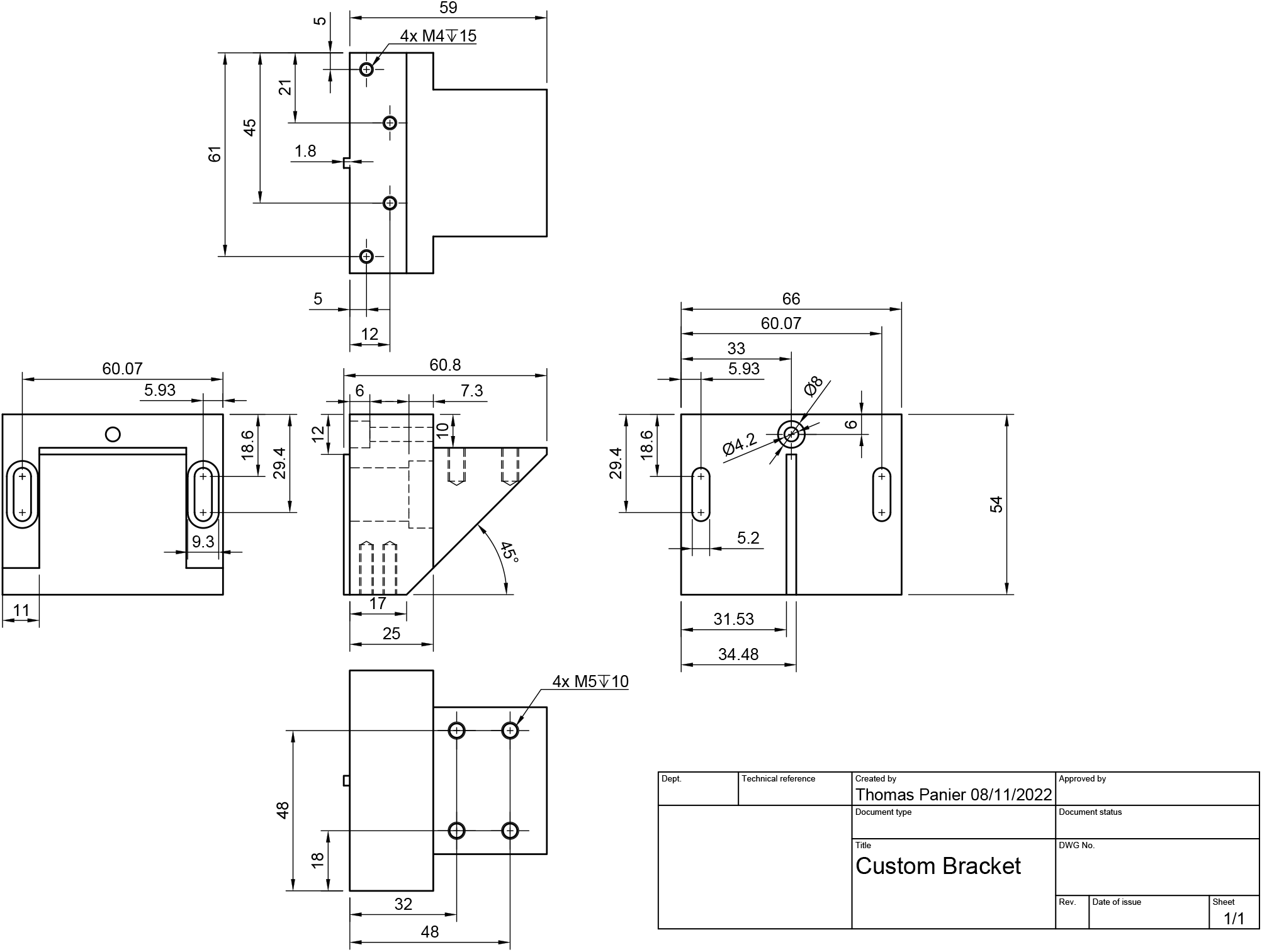
Custom bracket.

**Fig. S2.**
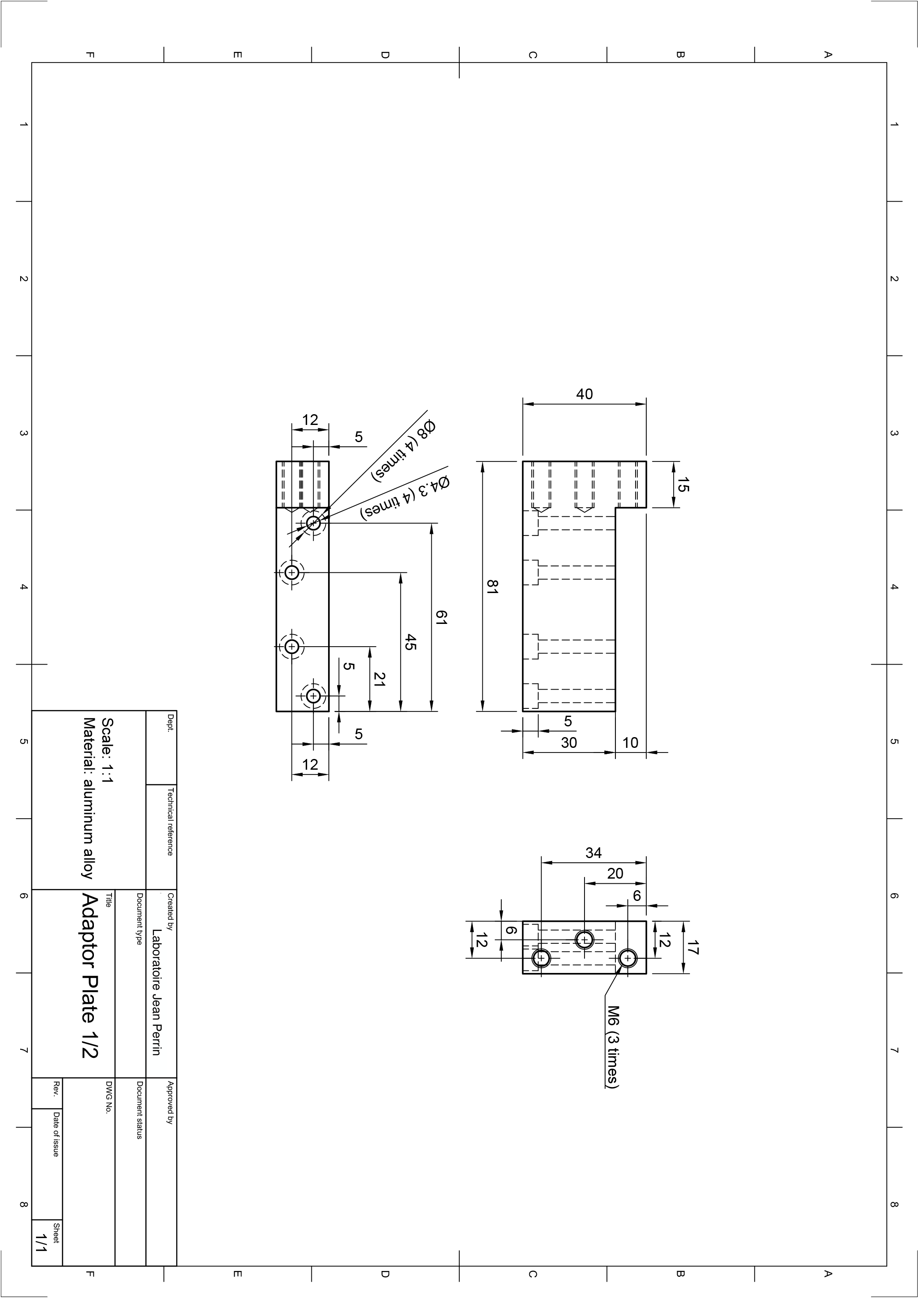
Adapter Plate 1.

**Fig. S3.**
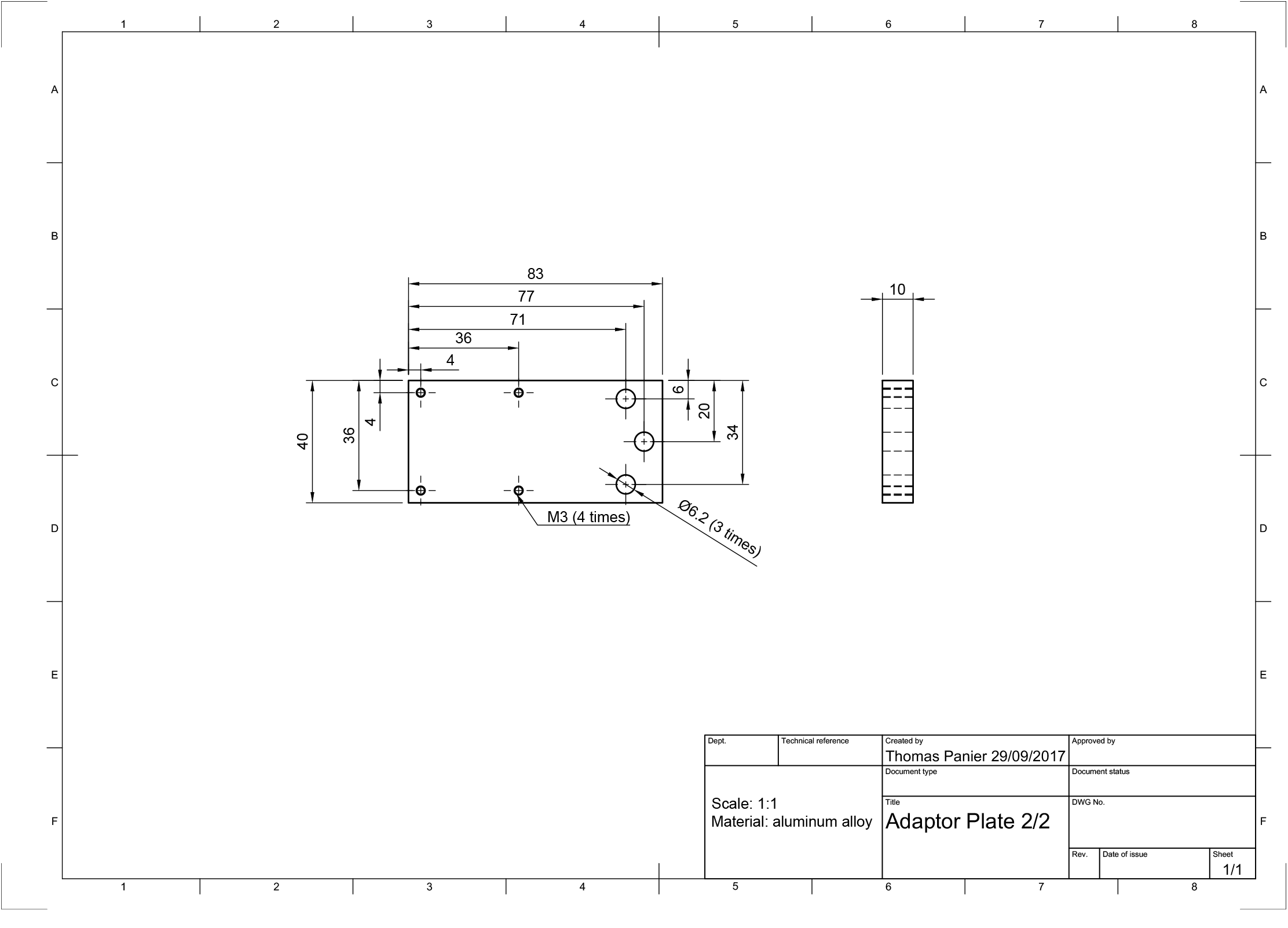
Adapter Plate 2.

**Fig. S4.**
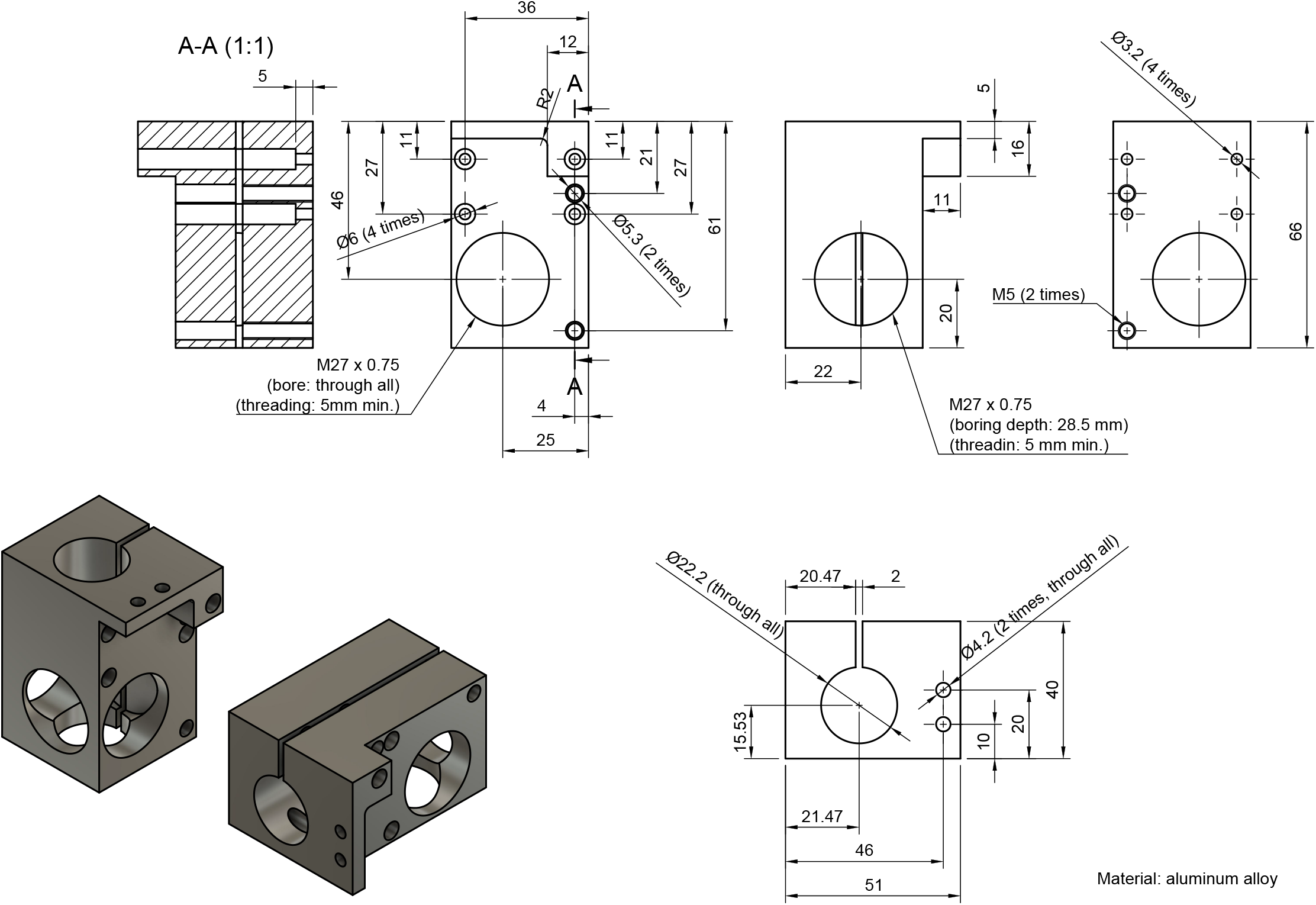
Light-sheet unit central cube.

**Fig. S5.**
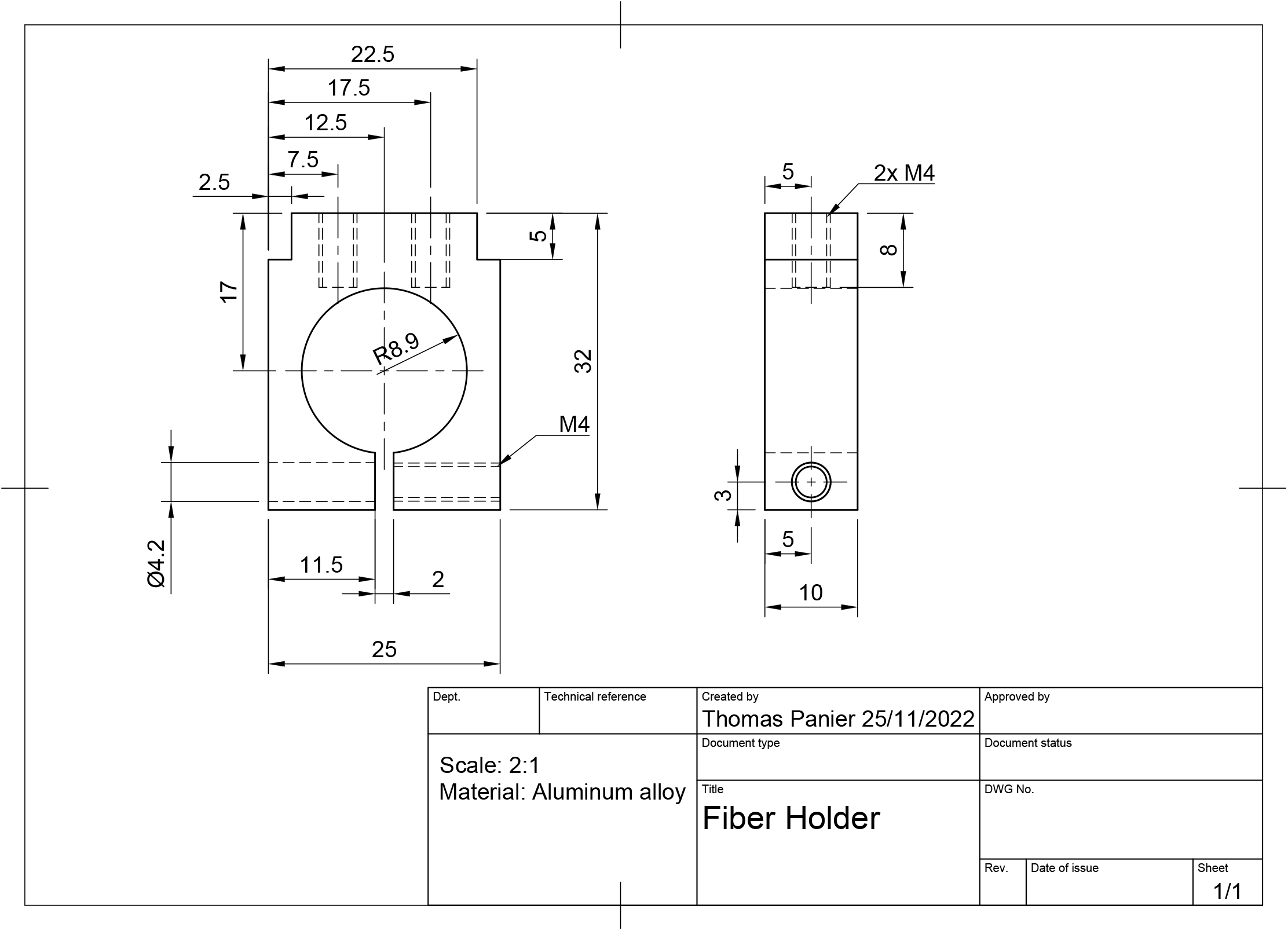
Fiber holder.

**Fig. S6.**
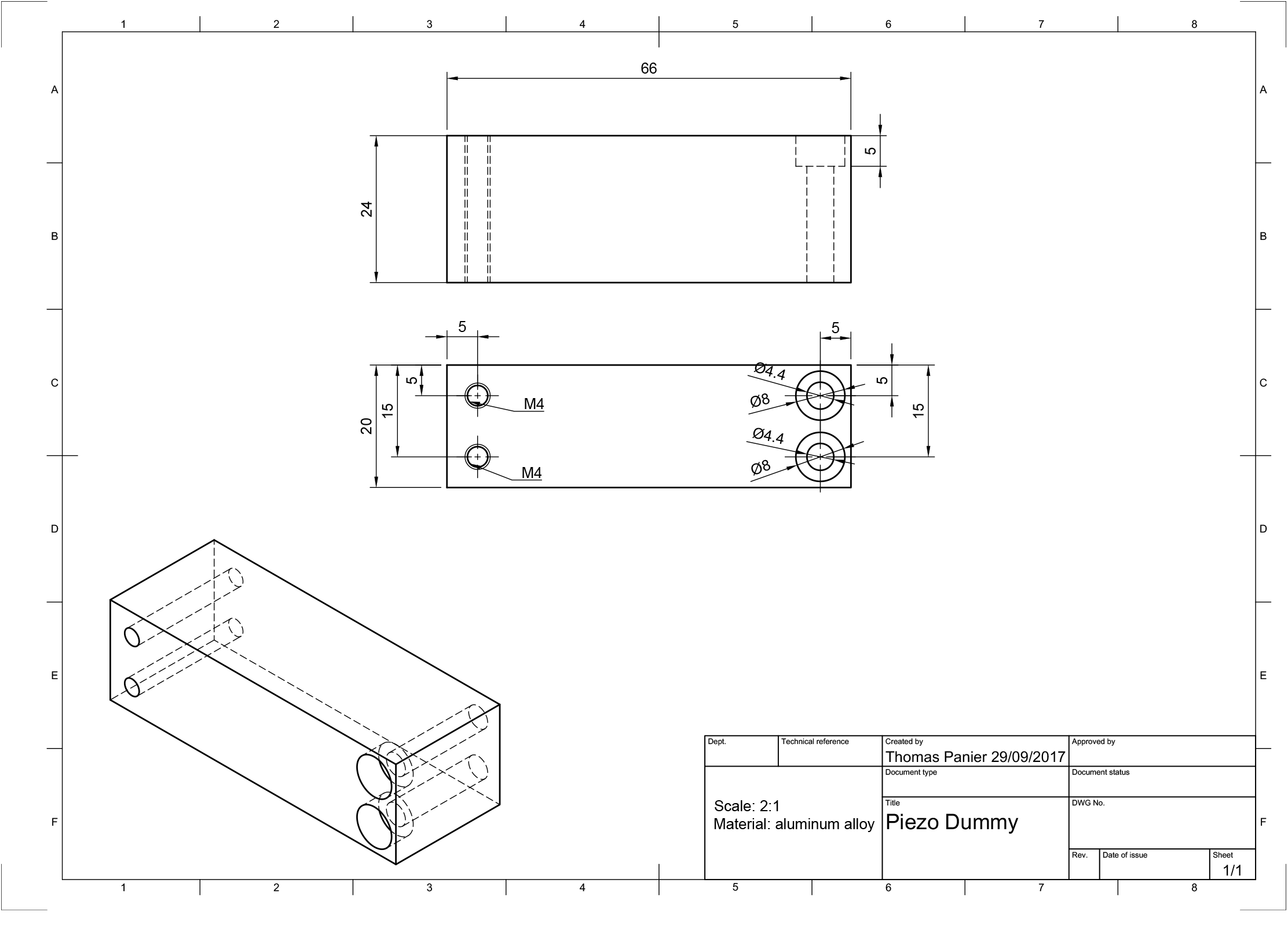
Piezo dummy.

**Fig. S7.**
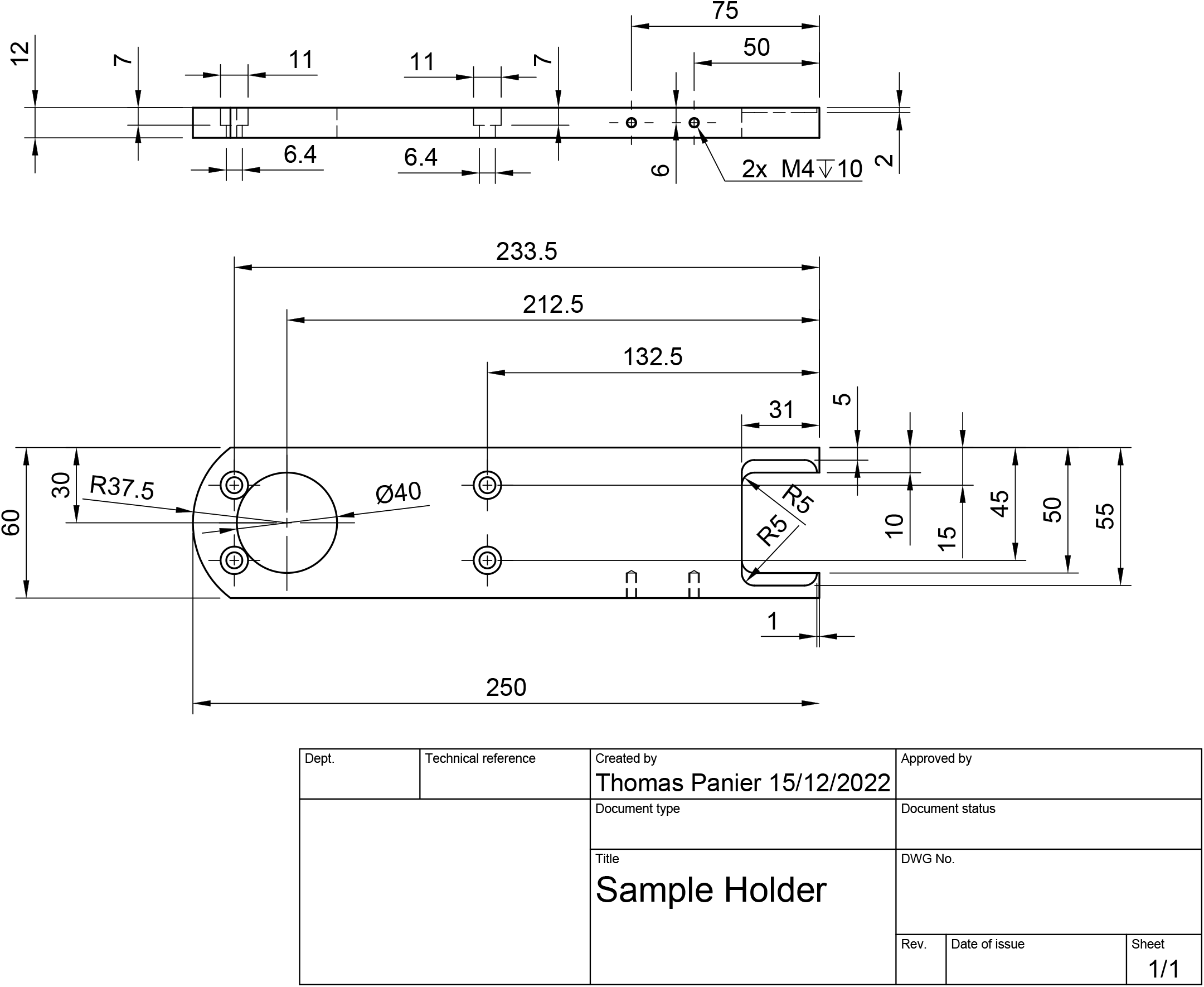
Sample holder.

**Fig. S8.**
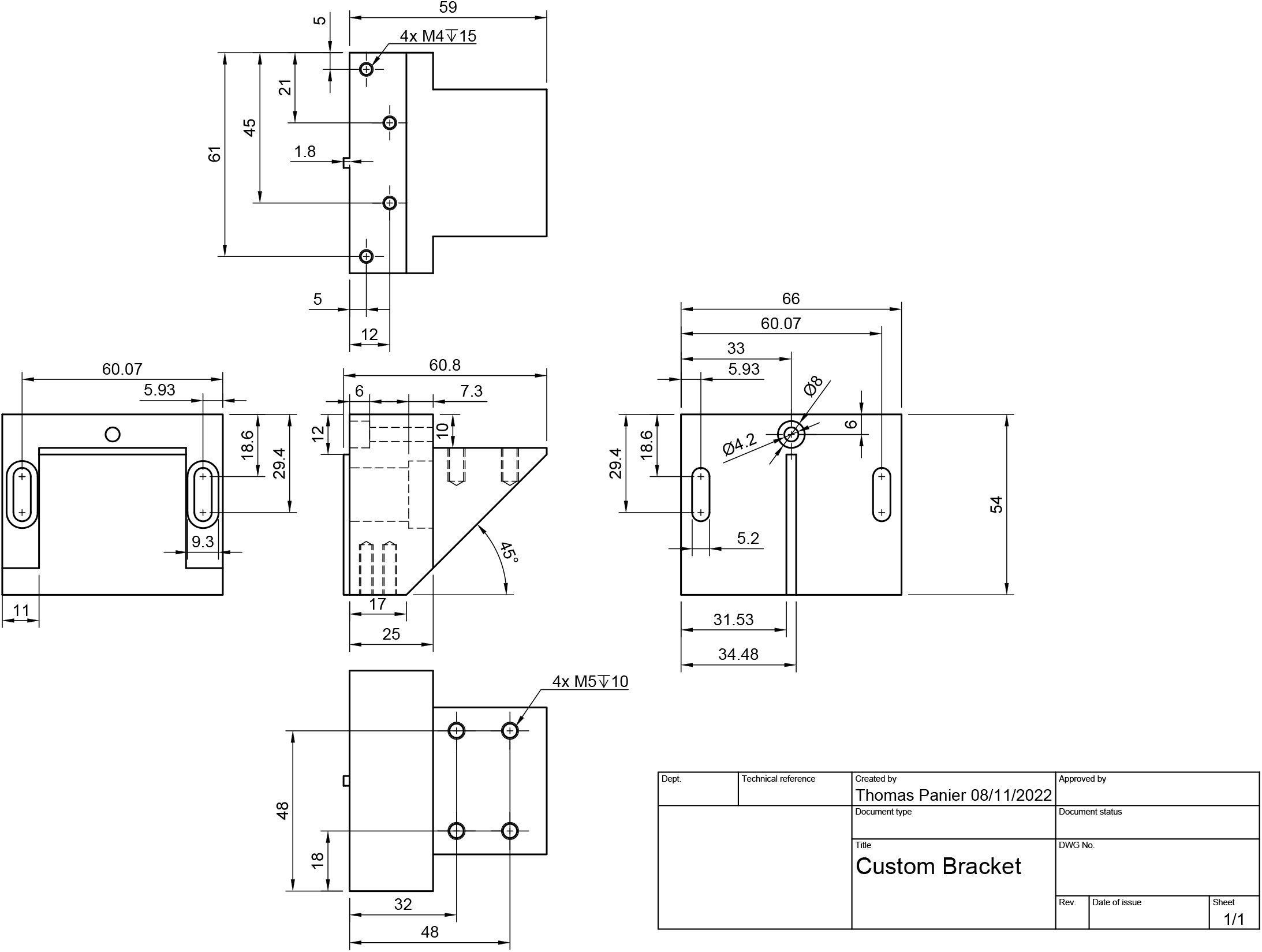
Sample holder.

